# Humanization of N-glycan-dependent protein quality control system in *Kluyveromyces marxianus* promotes glycoprotein secretion

**DOI:** 10.64898/2026.05.08.723133

**Authors:** Yi Ai, Yuting He, Lunqiang Zhao, Miaomiao Li, Yongming Wang, Jungang Zhou, Hong Lu, Yao Yu

**Affiliations:** State Key Laboratory of Genetics and Development of Complex Phenotypes, School of Life Sciences, Fudan University, Shanghai 200438, China; Shanghai Engineering Research Center of Industrial Microorganisms, Shanghai 200438, China

**Author notes:** Funding: National Key Research and Development Program of China (2021YFA0910603, 2021YFA0910601), Science and Technology Research Program of Shanghai (24HC2810100, 24ZR1406500, 2023ZX01).

**Keywords:** *Kluyveromyces marxianus*, ER glycoprotein folding, glycoprotein QC pathway humanization, recombinant glycoproteins

## Abstract

Human N-glycoproteins constitute a market worth hundreds of billions of dollars. However, their production in yeast is often limited by misfolding and subsequent degradation, largely due to differences in N-glycan-dependent protein quality control (QC) systems between humans and yeast. Notably, yeast lacks the UGGT-mediated reglucosylation-refolding cycle that rescues misfolded glycoproteins, and its degradation pathway involves fewer rate-limiting steps. To address this, we engineer the glycoprotein QC system in *Kluyveromyces marxianus*, a promising host for protein production, by introducing key human components and modifying native pathways. Expression of human UGGT1 or UGGT2 enhances the soluble and secretory production of glycoproteins in an activity-dependent manner. This effect is further improved by co-expression of the UGGT cochaperone SEP15 and by reducing native glucosidase II trimming activity. In addition, introduction of human EDEM2, a rate-limiting enzyme in glycoprotein degradation, delays ER-associated degradation and increases secretion. Integration of these engineering strategies substantially enhances the production of several high-value human-derived glycoprotein therapeutics, including etanercept, dulaglutide, and abatacept, with up to a ∼12-fold increase. These findings demonstrate that engineering a human-like glycoprotein QC network in yeast is an effective strategy to improve glycoprotein folding and secretion.

## 1. Introduction

N-glycosylation is one of the most prevalent post-translational modifications in eukaryotes and plays critical roles in protein folding, trafficking, and maturation^[1]^. The activity and stability of many therapeutic proteins depend on proper N-glycosylation. Nearly 40% of the world’s top-selling drugs, including monoclonal antibodies and Fc-fusion proteins, are recombinant N-glycoproteins derived from human proteins, collectively generating annual revenues exceeding USD 200 billion (https://www.pharmacircle.com/top_sales.php). These proteins are primarily produced in mammalian cell systems, such as CHO and SP2/0 cells, which possess a complete glycoprotein folding and processing machinery capable of generating human-like glycosylation structures^[2]^. Yeast expression systems, represented by *Saccharomyces cerevisiae* and *Komagataella phaffii* (*Pichia pastoris*), constitute another major platform for recombinant protein production and have enabled the commercial manufacture of products such as insulin, semaglutide, and the HPV vaccines, with annual sales exceeding USD 35 billion^[3]^. However, no commercially competitive human glycoprotein has yet been produced using yeast expression systems. This suggests that significant biological barriers still limit yeast as a mainstream platform for human glycoprotein production.

One major limitation is the structural disparity between yeast high-mannose N-glycans and the complex biantennary N-glycans typical of human glycoproteins^[4]^. This issue can be partially addressed through glycoengineering of the yeast glycosylation pathway^[5]^. A second and equally important challenge is the relatively low expression level of human glycoproteins in yeast. The highest reported titers of human antibodies and Fc-fusion proteins in yeast are approximately 1.6 g/L^[6]^ and 373 mg/L ^[7]^, respectively, which remain substantially lower than those achieved in CHO cells (∼10 g/L for antibodies^[8]^ and ∼5.5 g/L for Fc-fusion proteins^[9]^).

A key factor limiting the expression of human glycoproteins in yeast is the fundamental difference between yeast and human N-glycan-dependent protein quality-control systems. In humans, N-glycoproteins account for more than 21.3% of the proteome, compared with ∼6.1% in yeast^[10]^. Human cells therefore employ a stringent quality-control system that attempts repeated rescue of misfolded glycoproteins before degradation (**Figure 1**). In the endoplasmic reticulum (ER), nascent proteins receive the oligosaccharide Glc₃Man₉GlcNAc₂ (G3M9), which is sequentially trimmed by glucosidases I (GlsI) and II (GlsII) to produce Glc₁Man₉GlcNAc₂ (G1M9). This structure enters the calnexin/calreticulin (CNX/CRT) cycle, where lectin chaperones together with ER folding factors, including ERp57, ERp29 and cyclophilin B facilitate protein folding ^[11]^. During this process, GlsII removes the final glucose residue, generating Man₉GlcNAc₂ (G0M9). Correctly folded proteins are exported to the Golgi via the COPII machinery. If folding fails, UDP-glucose:glycoprotein glucosyltransferases (UGGT1/2) reglucosylate the glycan, with the aid of the cochaperone SEP15 ^[12]^, regenerating G1M9 and allowing re-entry into the CNX/CRT cycle for another round of folding. Persistent misfolding ultimately triggers sequential mannose trimming by EDEM2, followed by EDEM1 and EDEM3, leading to targeting for ER-associated degradation (ERAD)^[13]^.

**Figure 1.**
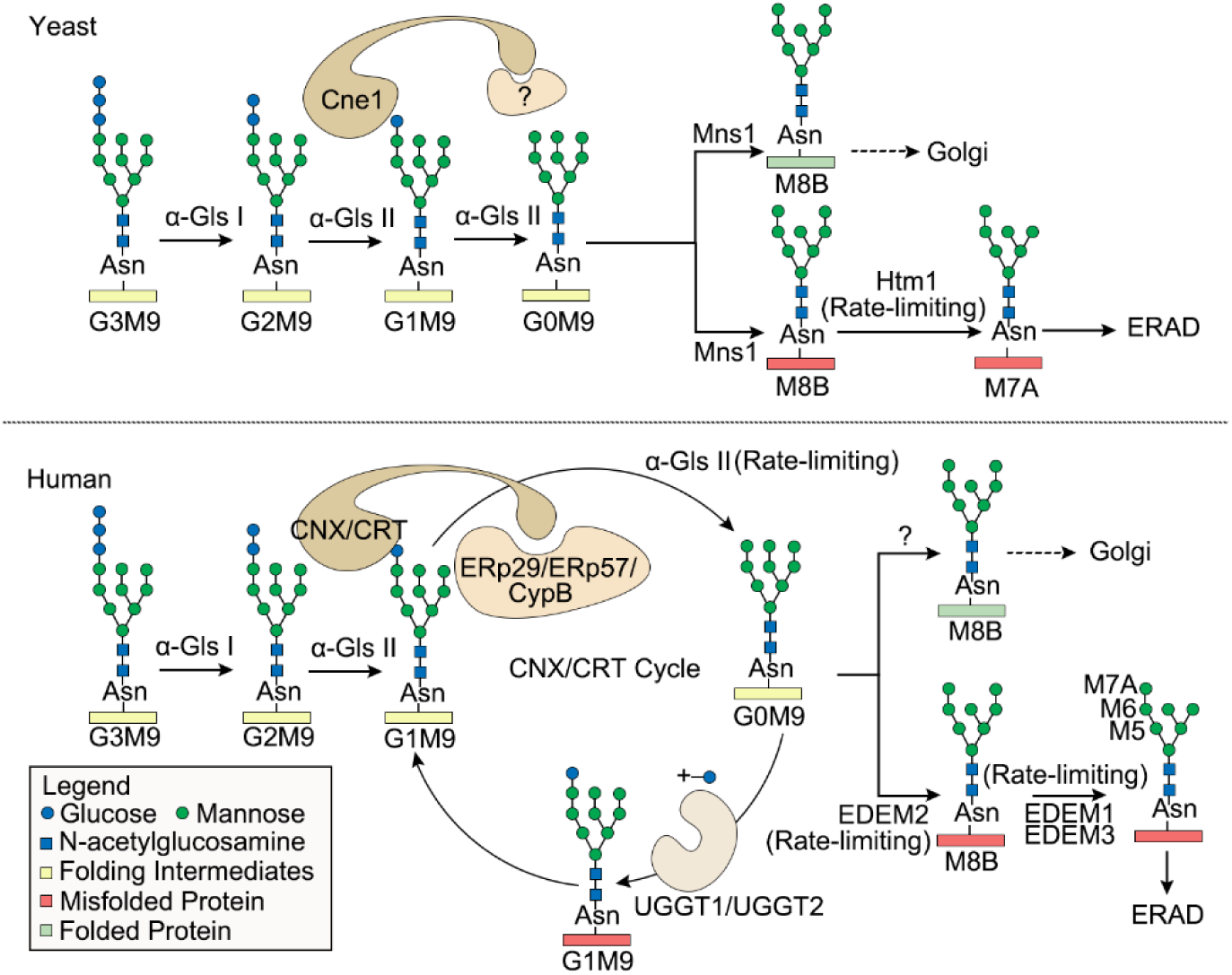
N-glycan-dependent quality control systems in yeast and human.

By contrast, most yeasts lack UGGT homologs and therefore cannot regenerate the monoglucosylated glycan required to re-enter the CNX/CRT cycle (Figure 1) ^[14]^. Moreover, key human cofactors such as ERp57 and ERp29 are absent in yeast, and the CNX/CRT homolog Cne1 lacks the C-terminal cytoplasmic tail of mammalian calnexin, which mediates various secondary functions, including calcium signaling^[10]^. Consequently, yeast glycoprotein quality control lacks the reglucosylation-based “refolding cycle” that characterizes human systems. In addition, the ERAD decision process is simplified in yeast. In human cells, sequential mannose trimming catalyzed by EDEM2 and EDEM3/EDEM1 constitutes two rate-limiting checkpoints for ERAD entry (Figure 1), whereas yeast relies primarily on a single rate-limiting enzyme, Htm1 (Figure 1)^[15]^.

Together, these differences indicate that yeast adopts a streamlined quality-control strategy for misfolded glycoproteins, which may cause premature degradation of heterologous human proteins and contribute to the low titers of human glycoproteins produced in yeast ^[16]^. Reconstituting aspects of the human glycoprotein quality-control system in yeast may provide a strategy to overcome this limitation. Consistent with this idea, co-expression of human calnexin in *S. cerevisiae* has been shown to improve the soluble expression of certain heterologous glycoproteins ^[17]^, indicating that components of the human glycoprotein quality-control machinery can function in yeast.

Here, we explored the effects of humanizing the glycoprotein quality-control system in *Kluyveromyces marxianus (K. marxianus)*. *K. marxianus* is an emerging food-grade yeast host capable of producing more than 50 heterologous proteins, including industrial enzymes, vaccines, and food proteins, with reported titers of up to 16.8 g/L^[18–23]^. Notably, *K. marxianus* produces relatively short N-glycan mannose chains (8–14 residues)^[22]^, similar to those of *P. pastoris*^[23]^ and substantially shorter than the hypermannosylated glycans (>50 residues) typical of *S. cerevisiae* ^[24]^. This feature makes *K. marxianus* a promising host for glycoengineering toward human glycoprotein production, and engineered strains capable of synthesizing Man₃GlcNAc₂/₄ precursors for complex human glycans have already been reported^[22]^.

In this study, we introduced key components of the human CNX/CRT cycle into *K. marxianus* and showed that UGGT1 and UGGT2 promote the folding and secretory expression of human glycoproteins in an enzyme activity–dependent manner. This effect was further enhanced by co-expression of SEP15 and by reducing GlsII trimming activity. Additionally, introduction of the human ERAD regulator EDEM2 delayed glycoprotein degradation and improved secretion. Combining these strategies resulted in up to a 12-fold increase in the production of various glycoproteins, including high-value human-derived therapeutics. Together, these results demonstrate that engineering a human-like glycoprotein quality-control network in yeast is an effective strategy to improve the folding, stability, and secretion of human glycoproteins.

## 2. Results

### 2.1. Human UGGT promotes soluble expression and secretion of glycoproteins in *K. marxianus* in a catalytic activity-dependent manner

In the mammalian ER, UGGT acts as a gatekeeper of glycoprotein quality control by recognizing misfolded substrates and reglucosylating their N-glycans, enabling re-entry into the calnexin/calreticulin folding cycle^[12]^. As *K. marxianus* lacks a UGGT homolog, we introduced human UGGT1 or its catalytic mutant UGGT1-D1358A ^[25]^, generating KM_UGGT1_ and KM_UGGT1-D1358A_. Owing to the large size (6800 bp) of the UGGT1 cassette, genome integration was performed using the high-efficiency Sta2-Cas9 system. Both proteins were successfully expressed (**Figure 2A**) without affecting cell growth (**Figure 2B**).

**Figure 2.**
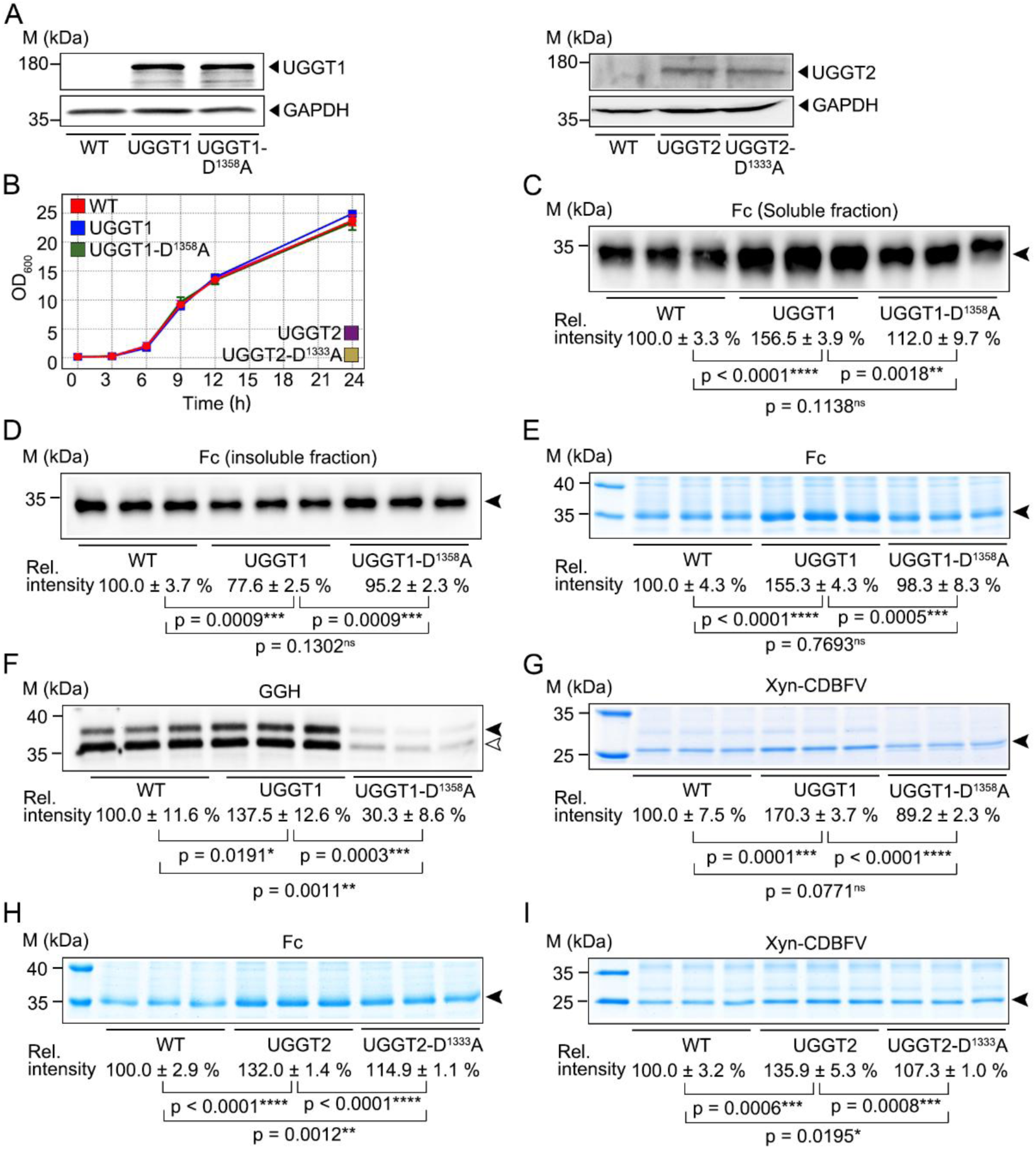
Effects of human UGGT1/UGGT2 and their catalytic mutants on glycoprotein secretion in *K. marxianus*. (A) Expression of human UGGT1, UGGT2, and their catalytic mutants in *K. marxianus*. WT, KM_UGGT1_ (UGGT1), KM_UGGT1-D1358A_ (UGGT1-D^1358^A), KM_UGGT2_ (UGGT2), and KM_U2-D1333A_ (UGGT2-D^1333^A) were cultured to exponential phase. Cell lysates were analyzed by WB using anti-UGGT1 or anti-UGGT2 antibodies. GAPDH served as a loading control. (B) Growth curves of WT and strains expressing UGGT1, UGGT2, or their mutants. Data are mean OD_600_ ±SD (n = 3). (C) Effect of UGGT1 on intracellular soluble Fc. The Fc plasmid (LHZ2068) was introduced into WT, KM_UGGT1_, or KM_UGGT1-D1358A_. After 72 h of shake-flask cultivation, cell lysates were analyzed by WB using an anti-His₆ antibody. No non-specific bands were detected at the Fc position in WT carrying the empty vector (**Figure S2**). (D) Effect of UGGT1 on intracellular insoluble Fc. Conditions were as in (C). Insoluble fractions were resuspended, boiled, and analyzed by WB. (E) Effect of UGGT1 on Fc secretion. After 24 h of bioreactor cultivation, supernatants were analyzed by SDS-PAGE. No bands were detected at the Fc position in WT carrying the empty vector (Figure S2). (F) Effect of UGGT1 on GGH secretion. The GGH plasmid (LHZ2069) was introduced, and supernatants were analyzed by WB after 72 h of shake-flask cultivation. Two bands were observed, with the smaller band (indicated by a white arrow) likely representing a degradation product. Neither band was detected in WT carrying the empty vector (Figure S2). (G) Effect of UGGT1 on Xyn-CDBFV secretion. The Xyn-CDBFV plasmid (LHZ443) was introduced, and supernatants were analyzed by SDS-PAGE after 72 h of shake-flask cultivation. No bands were detected at the Xyn-CDBFV position in WT carrying the empty vector (Figure S2). (H, I) Effects of UGGT2 on Fc (H) and Xyn-CDBFV (I) secretion. LHZ2068 or LHZ443 was introduced into KM_UGGT2_ or KM_U2-D1333A_, and analyses were performed as in (E) or (G), respectively. Black arrows indicate target proteins. Band intensities were quantified using ImageJ, with WT set to 100%. Data are mean ±SD (n = 3).

We first evaluated the human IgG Fc fragment, which contains a single N-glycosylation site. PNGase F treatment caused a clear mobility shift (**Figure S1**). Compared with WT, KM_UGGT1_ increased intracellular soluble Fc by 56%, reduced insoluble aggregates by 23%, and enhanced secretion by 55% (**Figure 2C–E**), indicating improved folding, solubility, and secretion. In contrast, UGGT1-D^1358^A showed no significant effect. To exclude the influence of glycosylation heterogeneity, supernatants were treated with PNGase F and the band at the expected molecular weight was quantified, yielding consistent results (**Figure S3**). These results indicate that UGGT1 promotes Fc production in a catalytic activity-dependent manner.

γ-Glutamyl hydrolase (GGH), a known UGGT substrate ^[26]^, was also glycosylated in *K. marxianus* (Figure S1). UGGT1 increased GGH secretion by 37%, whereas UGGT1-D^1358^A reduced it to ∼30% of WT (**Figure 2F**), possibly due to substrate trapping. Similar results were observed after PNGase F treatment (Figure S3), indicating that UGGT1 broadly enhances secretion of human glycoproteins.

We next examined the non-human glycoprotein xylanase Xyn-CDBFV ^[27]^. UGGT1 increased its secretion by 70%, whereas UGGT1-D^1358^A had no significant effect (**Figure 2G**). PNGase F-treated samples showed consistent results (Figure S3), demonstrating that UGGT1 promotes secretion of diverse glycoproteins in *K. marxianus*.

Finally, we evaluated the second isoform, UGGT2. The UGGT2 cassette (7258 bp) was integrated into the genome using Sta2-Cas9. Both UGGT2 and its catalytic mutant UGGT2-D^1333^A were expressed without affecting growth (Figure 2A-B). UGGT2 increased Fc secretion by 32%, whereas UGGT2-D^1333^A showed a reduced effect (15%) (**Figure 2H**). Similarly, UGGT2 enhanced Xyn-CDBFV secretion by 36%, while UGGT2-D^1333^A showed only a minor effect (7%) (**Figure 2I**). Thus, like UGGT1, UGGT2 promotes glycoprotein secretion in a catalytic activity-dependent manner, albeit less efficiently, likely due to differences in substrate specificity.

### 2.2. Cochaperone SEP15 enhances UGGT-mediated glycoprotein secretion

SEP15 is a cochaperone of UGGT that enhances its activity toward glycoprotein substrates^[12]^. To assess its effect, human SEP15 was introduced into *K. marxianus* strains expressing UGGT1 or UGGT2. The thioredoxin-like domain of SEP15 contains the rare amino acid selenocysteine (Sec), which is essential for its function ^[28]^. As *K. marxianus* lacks the Sec incorporation machinery, Sec was substituted with cysteine which retains comparable activity to wild-type SEP15 ^[25]^. SEP15 was successfully expressed in UGGT1- and UGGT2-containing strains (**Figure 3A and B**). Notably, native PAGE revealed an upward shift of UGGT1 and UGGT2 bands upon SEP15 co-expression, suggesting complex formation (Figure 3A and B).

**Figure 3.**
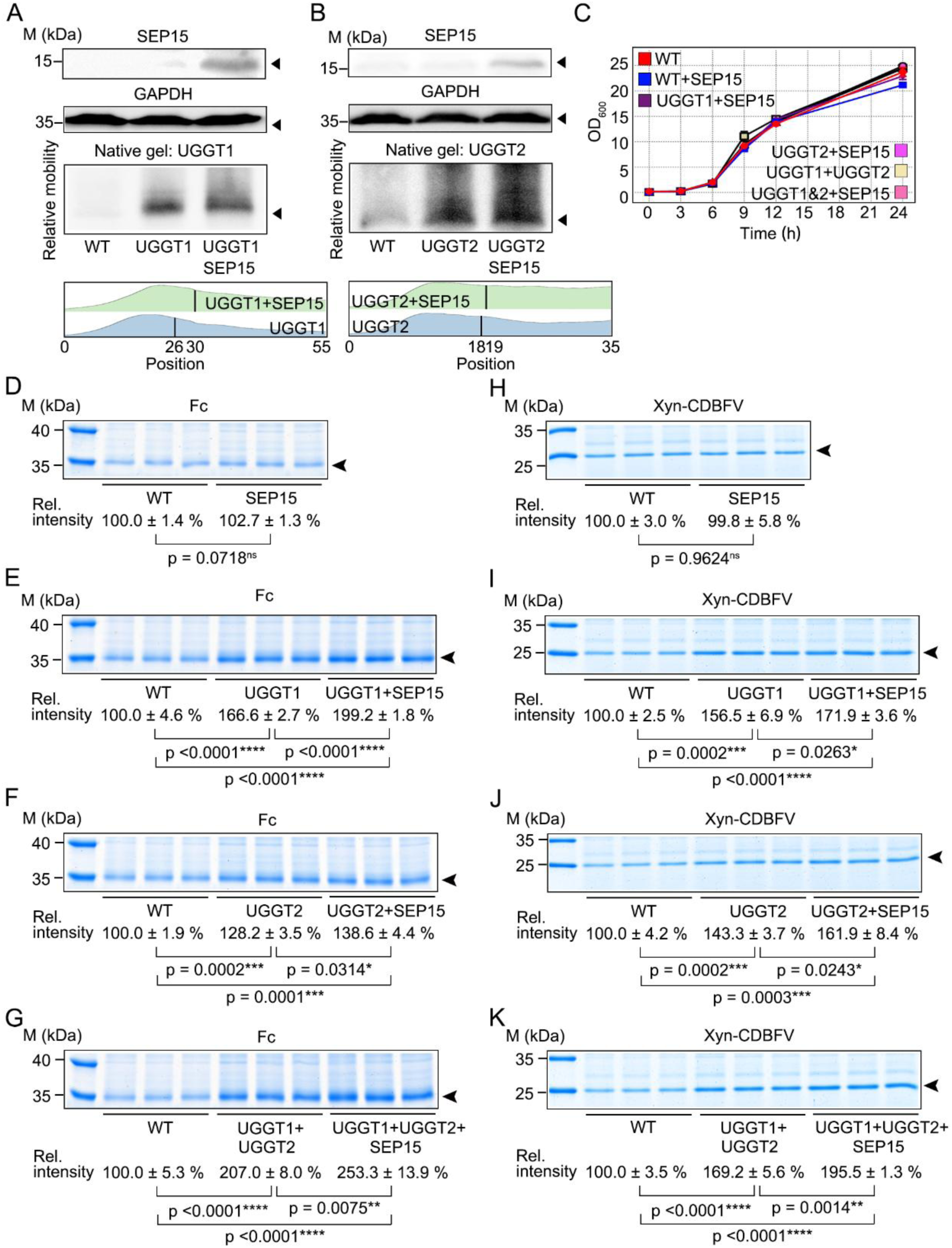
Cochaperone SEP15 enhances UGGT-mediated glycoprotein secretion. (A-B) SEP15 interacts with UGGT1 (A) and UGGT2 (B) in *K. marxianus*. WT, KM_UGGT1_ (UGGT1), KM_UGGT2_ (UGGT2), KM_U1-S_ (UGGT1 + SEP15), and KM_U2-S_ (UGGT2 + SEP15) were cultured to exponential phase. Cell lysates were analyzed by WB using SDS-PAGE (anti-SEP15 and anti-GAPDH) or native PAGE (anti-UGGT1 or anti-UGGT2). Black arrows indicate expected molecular weights. In native gels, grayscale intensity profiles along indicated lanes (bottom to top) are shown below the WB images, with medians indicated by vertical lines. (C) Growth curves of WT and SEP15-expressing strains. Data are mean OD_600_ ±SD (n = 3). (D–G) Effect of SEP15 on Fc secretion. The Fc plasmid (LHZ2068) was introduced into KM_SEP15_ (SEP15) (D), KM_U1-S_ (E), KM_U2-S_ (F), KM_U1&2-S_ (UGGT1 + UGGT2 + SEP15) (G), and control strains. After 24 h of bioreactor cultivation, supernatants were analyzed by SDS-PAGE. (H–K) Effect of SEP15 on Xyn-CDBFV secretion. The Xyn-CDBFV plasmid (LHZ443) was introduced into the indicated strains, and supernatants were analyzed by SDS-PAGE after 72 h of shake-flask cultivation. Black arrows indicate target proteins. Band intensities were quantified using ImageJ, with WT set to 100%. Data are mean ±SD (n = 3).

Expression of SEP15, either alone or together with UGGT1/UGGT2, had no effect on cell growth (**Figure 3C**). SEP15 alone did not enhance Fc secretion (**Figure 3D**). However, in UGGT1-expressing cells, SEP15 increased Fc secretion from 167% to 199% relative to WT (**Figure 3E**). In UGGT2-expressing cells, Fc secretion increased from 128% to 139% (**Figure 3F**). In strains co-expressing UGGT1 and UGGT2, SEP15 further increased Fc secretion from 207% to 253% (**Figure 3G**).

Similar trends were observed for Xyn-CDBFV. SEP15 alone had no effect, but in UGGT1-expressing cells, secretion increased from 157% to 172%; in UGGT2-expressing cells, from 143% to 162%; and in UGGT1/UGGT2 co-expression strains, from 169% to 196% (Figure 3H-K).

Together, these results demonstrate that SEP15 expression in *K. marxianus* further enhances UGGT1- and UGGT2-mediated secretion of diverse glycoproteins, likely through direct interaction that increases UGGT catalytic activity toward substrates.

### 2.3. Effects of human CRT/CNX and their cofactors on glycoprotein secretion in *K. marxianus*

In addition to UGGT and SEP15, we examined whether other components of the human CNX/CRT cycle could enhance glycoprotein secretion. *K. marxianus* contains a CNX/CRT homolog, Cne1. To avoid potential interference, we first evaluated its role. Deletion of *CNE1* had no effect on growth (Figure 4A) but caused a modest yet significant decrease in Fc secretion compared with WT (Figure 4B), indicating that endogenous Cne1 partially supports glycoprotein secretion.

**Figure 4.**
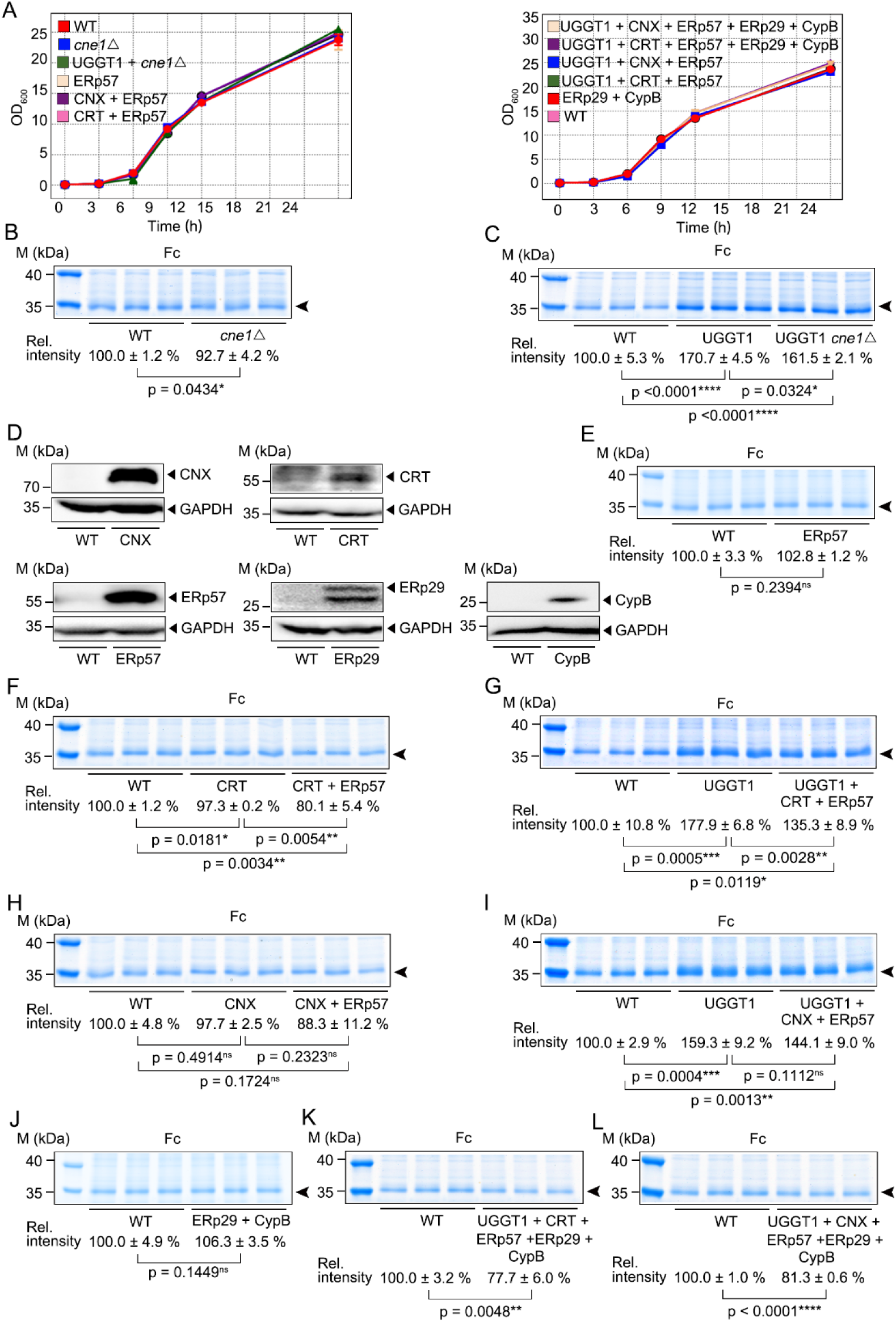
Effects of human CRT/CNX and their cofactors on glycoprotein secretion in *K. marxianus*. (A) Growth curves of WT, *cne1*Δ mutant and strains expressing human CRT/CNX cycle components. Data are mean OD_600_ ± SD (n = 3). (B-C) Effect of deleting *CNE1* on the UGGT1-mediated Fc secretion. The Fc plasmid (LHZ2068) was introduced into WT, KM_cne1_(*cne1*Δ) (B), KM_U1-*cne1*Δ_ (UGGT1+*cne1*Δ) (C). After 24 h of bioreactor cultivation, supernatants were analyzed by SDS-PAGE. (D) Expression of human CRT/CNX cycle components in *K. marxianus*. WT, KM_CNX_ (CNX), KM_CRT_ (CRT), KM_p57_ (ERp57), KM_cypb-p29_ (ERp29), KM_cypb-p29_ (CypB) were cultured to exponential phase. Cell lysates were analyzed by WB using primary antibodies against human components. GAPDH served as a loading control. (E-L) Effects of human CRT/CNX cycle components on UGGT1-mediated Fc secretion. LHZ2068 was introduced into KM_p57_ (ERp57) (E), KM_CRT_(CRT) (F), KM_CRT-p57_(CRT+ERp57) (F), KM_U1-CRT-p57_ (UGGT1+CRT+ ERp57) (G), KM_CNX_(CNX) (H), KM_CNX-p57_(CNX+ERp57) (H), KM_U1-CNX-p57_ (UGGT1+CNX+ERp57) (I), KM_cypb-p29_ (CypB+ERp29) (J), KM_U1-CRT-3C_ (UGGT1+CRT+ERp57+ERp29+CypB) (K),KM_U1-CNX-3C_ (UGGT1+CNX+ERp57+ERp29+CypB) (L). Cultivation and analyses were performed as in (B). Arrows indicate Fc bands. Band intensities were quantified using ImageJ, with WT set to 100%. Data are mean ±SD (n = 3).

Notably, deletion of CNE1 in the UGGT1 background reduced Fc secretion by ∼9% relative to UGGT1 alone (**Figure 4C**), similar to the reduction observed in the WT background (∼7%). This additive effect suggests that Cne1 and UGGT1 act through distinct pathways, consistent with reports that yeast Cne1 does not participate in a canonical CNX/CRT cycle but instead functions in other processes, such as β-1,6-glucan synthesis ^[29]^.

Given this, we introduced human CNX, CRT, and their cofactors (ERp57, ERp29, and CypB) to assess their effects. All proteins were expressed without affecting growth (Figure 4A and D). Expression of ERp57 or CRT alone had minimal impact on Fc secretion (Figure 4E, F). However, co-expression of CRT and ERp57 reduced Fc secretion by ∼20% relative to WT, suggesting that their interaction impairs folding and secretion (Figure 4F). This negative effect was more pronounced in the UGGT1 background, where Fc secretion decreased from 178% to 135% upon co-expression of CRT and ERp57 (Figure 4G). Similarly, co-expression of CNX and ERp57 slightly reduced Fc secretion in both WT and UGGT1 strains, although the effect was weaker and not significant (Figure 4H, I).

Co-expression of ERp29 and CypB in WT had no effect on Fc secretion (Figure 4J). However, in the UGGT1+CRT+ERp57 or UGGT1+CNX+ERp57 backgrounds, co-expression of ERp29 and CypB markedly reduced Fc secretion (from 135% to 77% and from 144% to 81%, respectively) (Figure 4K, L). These results indicate that ERp29 and CypB interfere with glycoprotein folding, not only abolishing the UGGT1-mediated enhancement of Fc secretion but also reducing secretion below WT levels.

The above results suggest that, under the current conditions, CNX/CRT and their cofactors (ERp57, ERp29, and CypB) do not form an effective CNX/CRT cycle with UGGT in *K. marxianus* to enhance Fc production. Instead, they likely compete with and interfere with UGGT-mediated glycoprotein folding.

### 2.4. Engineering the GlsII β-subunit improves glycoprotein secretion in *K. marxianus*

In humans, GlsII trims G2M9 glycans to G1M9 and G0M9, and G0M9 can be reglucosylated by UGGT to regenerate G1M9, forming the basis of the reglucosylation–refolding cycle. To assess functional differences between human and yeast GlsII, we deleted the endogenous GlsII α- and β-subunit genes (*ROT2* and *GTB1*) in *K. marxianus* to generate KM_GIIΔ_, and replaced them with the human α- and β-subunits (*GANAB* and *PRKCSH*) to generate KM_hGII_. GANAB and PRKCSH were successfully expressed (**Figure 5A**), and neither strain showed growth defects (**Figure 5B**).

**Figure 5.**
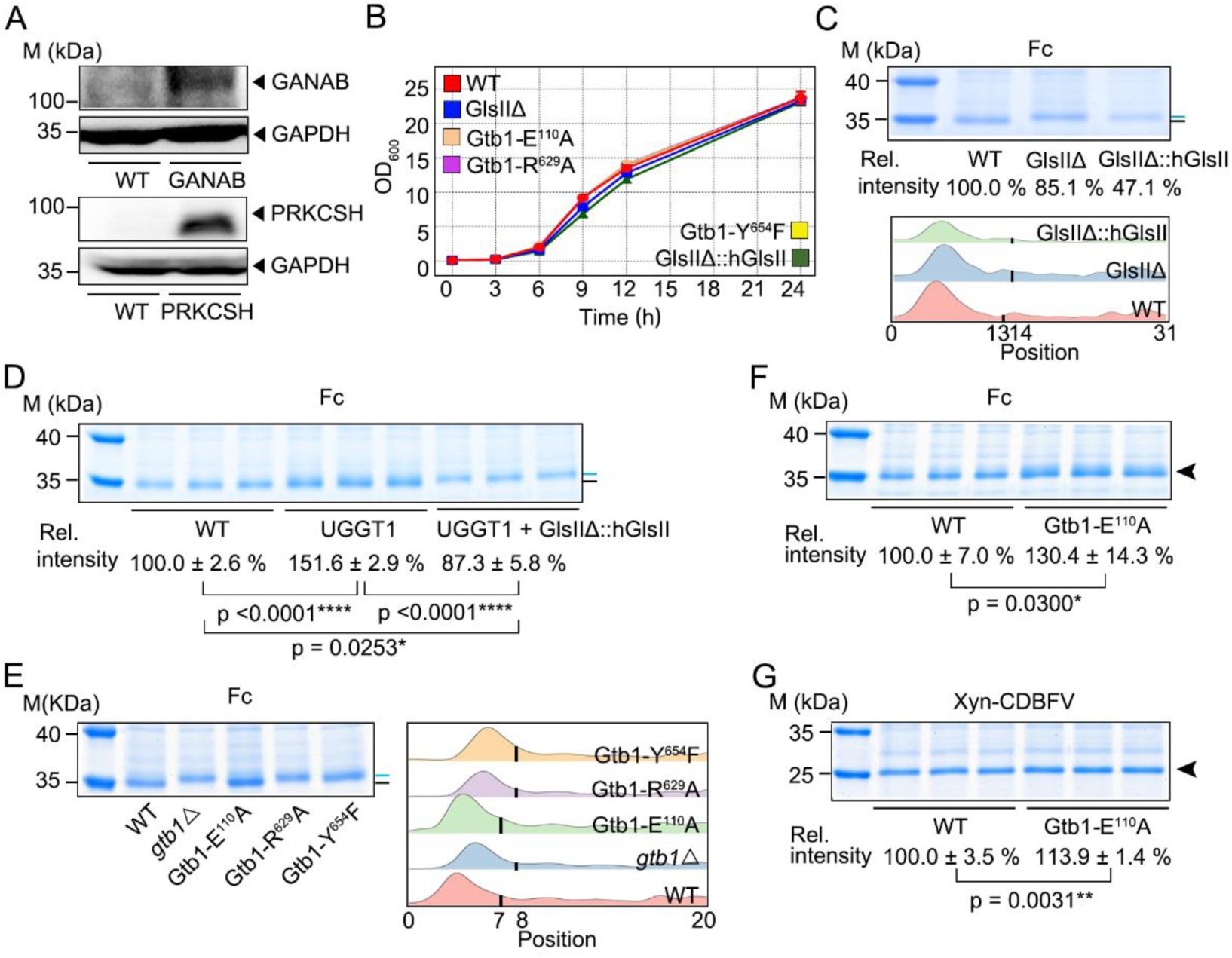
Engineering the GlsII β-subunit improves glycoprotein secretion in *K. marxianus*. **(A)** Expression of human GANAB and PRKCSH in *K. marxianus*. WT and KM_hGII_ strains expressing GANAB and PRKCSH were cultured to exponential phase. Cell lysates were analyzed by WB using anti-GANAB or anti-PRKCSH antibodies. GAPDH served as a loading control. **(B)** Growth curves of WT, the GlsII deletion mutant, strains expressing human GlsII, and the GTB1 point mutant. WT, KM_GIIΔ_ (GlsIIΔ), KM_hGII_ (GlsIIΔ::hGlsII), KM_β-E110A_ (Gtb1-E^110^A), KM_β-R629A_ (Gtb1-R^629^A), and KM_β-Y654F_ (Gtb1-Y^654^F) were cultured in YPD for 24 h. Data are mean OD_600_ ±SD (n = 3). **(C)** Effect of GlsII mutants on Fc mobility shift and secretion. The Fc plasmid (LHZ2068) was introduced into WT, KM_GIIΔ_, and KM_hGII_. After 24 h of bioreactor cultivation, supernatants were analyzed by SDS-PAGE. One representative gel is shown, and an additional replicate is provided in Figure S4. Grayscale intensity profiles along indicated lanes (bottom to top) are shown below the gel, with medians indicated by vertical lines. **(D)** Effect of replacing native GlsII with human counterparts on UGGT1-mediated Fc secretion. LHZ2068 was introduced into WT, KM_U1_ (UGGT1), and KM_U1-hGII_ (UGGT1 + GlsIIΔ::hGlsII). **(E, F)** Effects of *GTB1* mutations on Fc mobility shift (E) and secretion (F). LHZ2068 was introduced into KM_gtb1Δ_(*gtb1*Δ), KM_β-E110A_, KM_β-R629A_, and KM_β-Y654F_. **(G)** Effect of the Gtb1-E^110^A mutation on Xyn-CDBFV secretion. The Xyn-CDBFV plasmid (LHZ443) was introduced into the indicated strains, and supernatants were analyzed by SDS-PAGE after 72 h of shake-flask cultivation. Cultivation and SDS-PAGE analyses in (D–F) were performed as in (C). Band-shift analysis in (E) was performed as in (C). In (D–F), blue and black lines indicate Fc bands with or without mobility shift, respectively. In (G), the black arrow indicates Xyn-CDBFV. Band intensities in (C–G) were quantified using ImageJ, with WT set to 100%. Data in (C) are mean (n = 2), and in (D–G), mean ±SD (n = 3).

Expression of Fc in KM_GIIΔ_ reduced secretion compared with WT, confirming that GlsII supports glycoprotein production^[30]^. Fc also showed a higher molecular weight in KM_GIIΔ_ (**Figure 5C**), consistent with impaired trimming of G2M9/G1M9 glycans and retention of glucose residues ^[31]^. A similar shift was observed in KM_hGII_ (Figure 5C), indicating that human GlsII could not functionally replace the yeast enzyme. Accordingly, Fc secretion in KM_hGII_ was not restored and was even lower than in KM_GIIΔ_ (Figure 5C), suggesting that human GlsII disrupts glycoprotein processing.

In the UGGT1 background, replacement with human GlsII (KM_U1-hGII_) abolished the UGGT1-mediated enhancement of Fc secretion (**Figure 5D**), indicating that UGGT1 activity depends on functional GlsII to generate G0M9 substrates.

Because heterologous human GlsII failed to improve secretion, we hypothesized that tuning native GlsII activity to slow G2M9/G1M9 trimming could prolong folding time and reduce ERAD. Gtb1, the regulatory subunit of GlsII, was targeted for engineering. Based on conserved residues implicated in trimming efficiency in *S. cerevisiae* ^[31]^, we generated Gtb1-E^110^A, -R^629^A, and -Y^654^F mutants (Figure S5), none of which showed growth defects (Figure 5B).

Fc expressed in Gtb1-R^629^A and Gtb1-Y^654^F showed pronounced upward mobility shifts, comparable to the *gtb1*Δ mutant, indicating severe loss of GlsII function and accumulation of incompletely trimmed glycans. In contrast, Gtb1-E^110^A showed WT-like mobility, indicating retained activity (**Figure 5E**). Notably, Gtb1-E^110^A increased Fc secretion by ∼30% (**Figure 5F**) and Xyn-CDBFV secretion by ∼14% (**Figure 5G**), likely due to moderately slowed glycan trimming that extends folding time.

### 2.5. Human EDEM2 delays glycoprotein degradation and enhances secretion in *K. marxianus*

In humans, G0M9 on misfolded glycoproteins is trimmed by EDEM2 to M8B, representing the first rate-limiting step for ERAD entry, followed by EDEM3 (primarily) and EDEM1 (secondarily) processing M8B to M7A as a second rate-limiting step (Figure 1)^[15]^. In contrast, yeast Mns1-mediated G0M9-to-M8B trimming is not rate-limiting. Instead, Htm1-catalyzed M8B-to-M7A conversion serves as the sole rate-limiting step for ERAD entry^[32]^.

To mimic this more stringent ERAD entry control, we introduced human EDEM2 or its catalytic mutant EDEM2-E^117^Q into *K. marxianus* ^[15]^. Both proteins were expressed (Figure 6A) without affecting growth (Figure 6B). Upon cycloheximide treatment, Fc degradation in the EDEM2 strain was faster than in *mns1*Δ strain but slower than in WT, indicating that EDEM2 delays ERAD but does not severely impair it, as observed in *mns1*Δ strain (Figure 6C). In contrast, Fc degradation in the EDEM2-E^117^Q strain was comparable to WT, demonstrating that this effect depends on EDEM2 mannosidase activity (Figure 6C). Consistently, EDEM2 increased Fc secretion by 52%, whereas EDEM2-E^117^Q showed no improvement (Figure 6D). Notably, *mns1*Δ strain also enhanced Fc secretion (∼36%), but to a lesser extent than EDEM2 (Figure 6E).

**Figure 6.**
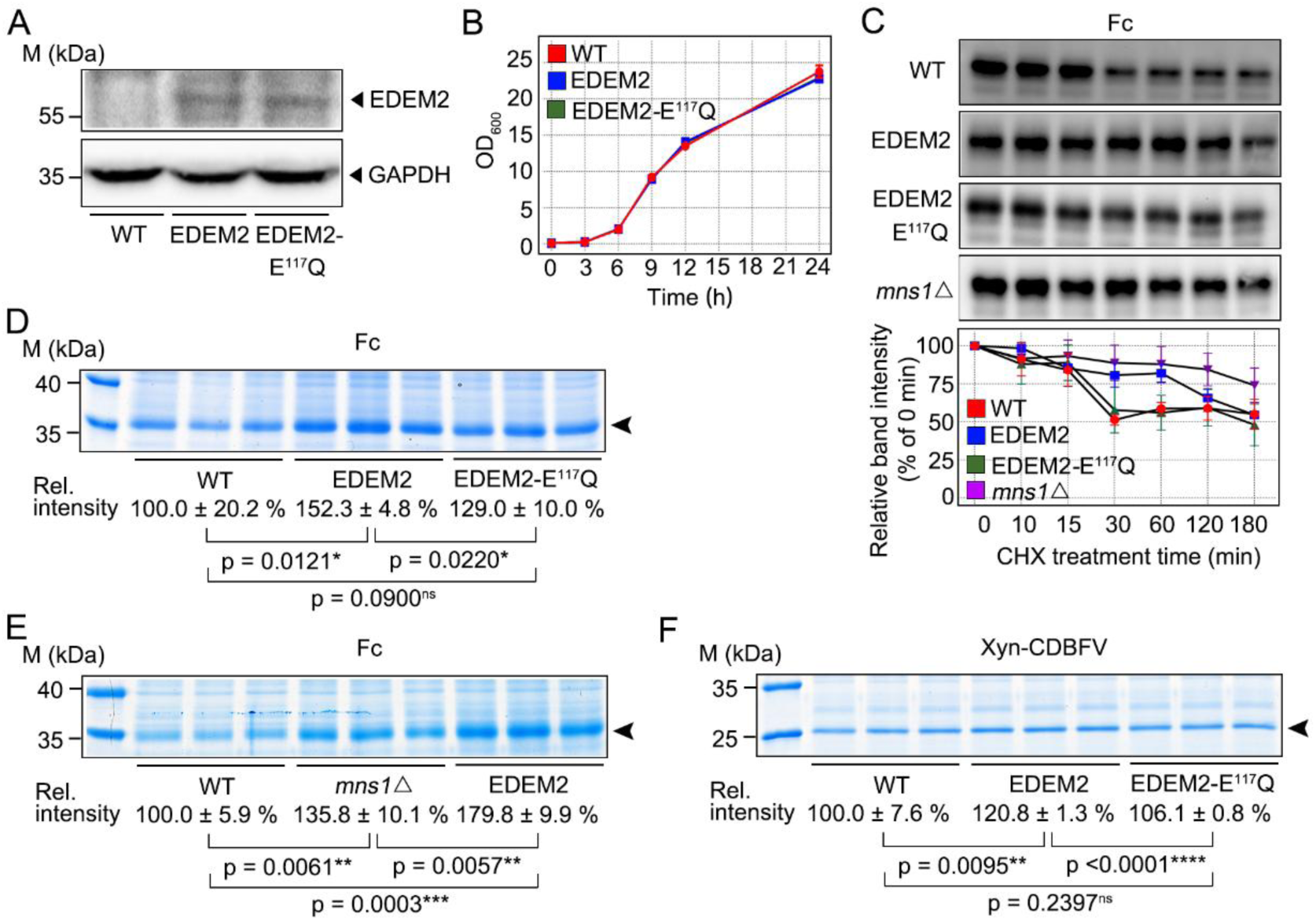
Human EDEM2 delays glycoprotein degradation and enhances secretion in *K. marxianus*. (A) Expression of human EDEM2 and its catalytic mutant in *K. marxianus*. WT, KM_EDEM2_ (EDEM2), and KM_E-E117Q_ (EDEM2-E^117^Q) were cultured to exponential phase. Cell lysates were analyzed by WB using an anti-EDEM2 antibody, with GAPDH as a loading control. (B) Growth curves of WT and EDEM2-expressing strains. Data are mean OD_600_ ±SD (n = 3). (C) Effect of EDEM2 on Fc degradation. The Fc plasmid (LHZ2068) was introduced into WT, KM_EDEM2_, KM_E-E117Q_, and KM*_mns1_*_Δ_ (*mns1*Δ). After 24 h of shake-flask cultivation, cells were treated with cycloheximide (CHX) for the indicated times. Cell lysates were analyzed by WB using an anti-His₆ antibody. Representative images are shown (top), and additional replicates are provided in Figure S6. Fc band intensity at 0 min was set to 100% for normalization, and relative protein levels over time are shown (bottom). Data are mean ±SD (n = 3). **(D-E)** Effects of EDEM2 (D) and *MNS1* deletion (E) on Fc secretion. Supernatants from 24 h bioreactor cultures were analyzed by SDS-PAGE. **(F)** Effect of EDEM2 on Xyn-CDBFV secretion. The Xyn-CDBFV plasmid (LHZ443) was introduced into WT, KM_EDEM2_, and KM_E-E117Q_. Supernatants were analyzed by SDS-PAGE after 72 h of shake-flask cultivation. In (D-F), black arrows indicate target proteins. Band intensities were quantified using ImageJ, with WT set to 100%. Data are mean ±SD (n = 3).

A similar trend was observed for Xyn-CDBFV, where EDEM2 increased secretion by 20%, while EDEM2-E^117^Q had no effect (**Figure 6F**). These results indicate that EDEM2 promotes glycoprotein secretion by delaying degradation in an activity-dependent manner, and is more effective than severely impairing ERAD (e.g., *MNS1* deletion).

### 2.6. Combined engineering of the glycoprotein quality control pathway significantly enhances secretion

We combined effective glycoprotein quality control engineering strategies to further improve secretion (**Figure 7A**). In the UGGT1/2 + SEP15 background, we introduced either the Gtb1-E^110^A or human EDEM2. Growth curves showed no apparent differences from WT within 24 h (**Figure 7B**). Introduction of Gtb1-E^110^A increased Fc secretion from 247% to 285% (**Figure 7C**), while EDEM2 increased secretion from 257% to 289% (**Figure 7D**). These results indicate that, in the presence of UGGT-mediated reglucosylation, either slowing GlsII trimming or delaying ERAD entry further enhances glycoprotein secretion in *K. marxianus*.

**Figure 7.**
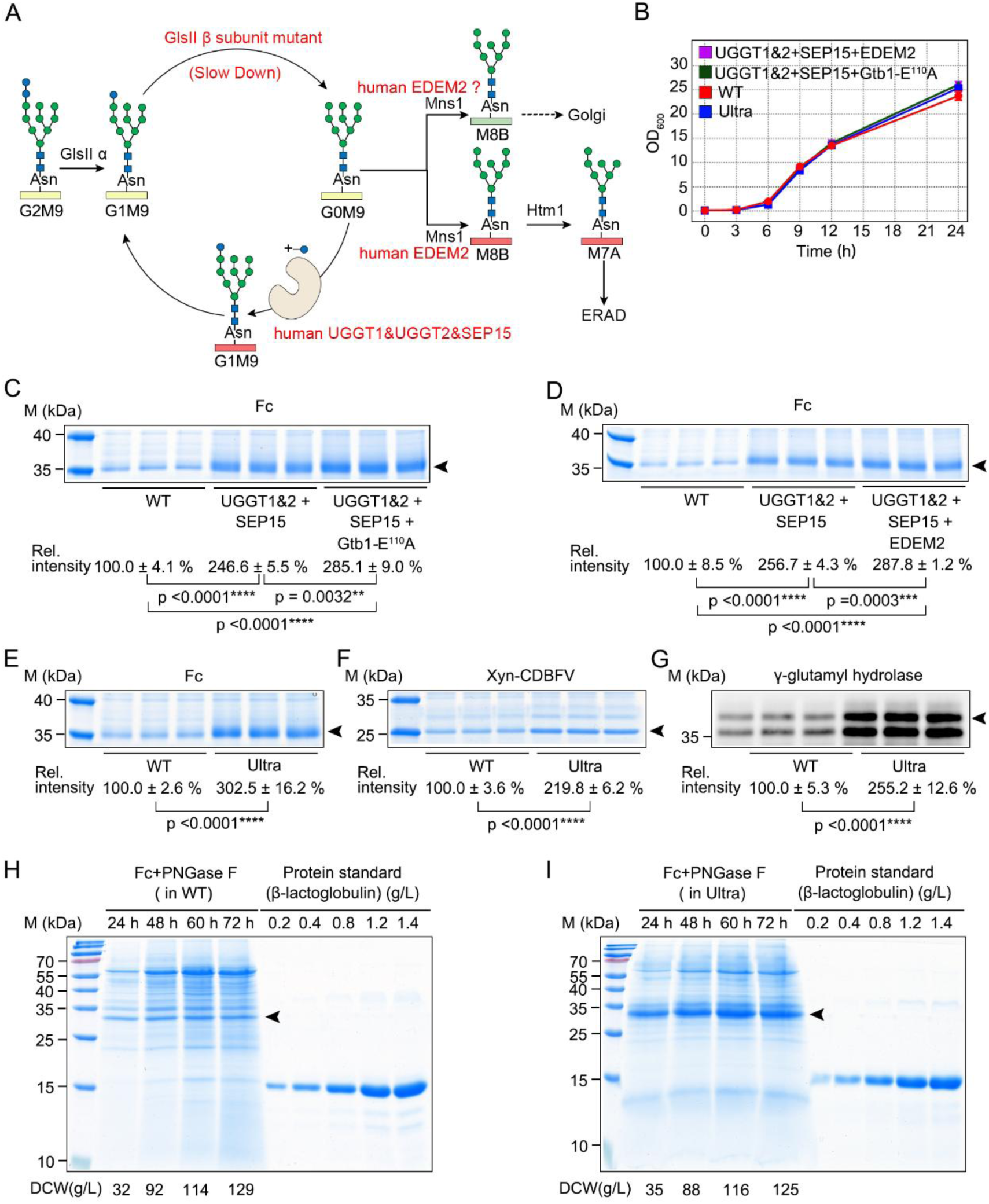
Combined engineering of the glycoprotein quality control pathway significantly enhances secretion. **(A)** Schematic overview of glycoprotein quality control pathway engineering. **(B)** Growth curves of WT, KM_U1&2-S-E110A_ (UGGT1&2 + SEP15 + Gtb1 E^110^A), KM_U1&2-S-EDEM2_ (UGGT1&2 + SEP15 + EDEM2), and KM_ultra_ (Ultra). Data are mean OD_600_ ±SD (n = 3). **(C, D)** Gtb1-E^110^A mutation (C) and EDEM2 (D) enhance UGGT1&2-SEP15-mediated secretion. The Fc plasmid (LHZ2068) was introduced into WT, KM_U1&2-S_, KM_U1&2-S-E110A_, and KM_U1&2-S-EDEM2_. Supernatants were analyzed by SDS-PAGE after 24 h bioreactor cultivation. **(E–G)** Combined engineering of five modifications further improves secretion of Fc (E), Xyn-CDBFV (F), and GGH (G). LHZ2068 (Fc), LHZ2069 (GGH), and LHZ443 (Xyn-CDBFV) plasmids were introduced into WT and KM_ultra_. Fc secretion was analyzed as in (C), while GGH was analyzed by WB using an anti-His₆ antibody, Xyn-CDBFV was analyzed by SDS-PAGE after 72 h shake-flask cultivation. In (C–G), arrows indicate target proteins. Band intensities were quantified using ImageJ, with WT set to 100%. Data are mean ±SD (n = 3). (**H-I**) Fc titer in WT and KM_ultra_. Supernatants were collected after 24, 48, 60, and 72 h of bioreactor cultivation and analyzed by SDS-PAGE following PNGase F treatment. SDS-PAGE of untreated supernatants is shown in Figure S7. Quantification was performed using β-lactoglobulin standards, and the calibration curve is shown in Figure S8.

We then combined all effective modifications-UGGT1, UGGT2, SEP15, Gtb1-E^110^A, and EDEM2-to generate the KM_ultra_ strain (also called KM-AY). KM_ultra_ exhibited growth comparable to WT (Figure 7B). It increased Fc secretion to 302% of WT (**Figure 7E**), Xyn to 220% (**Figure 7F**), and γ-glutamyl hydrolase to 255% (**Figure 7G**), representing the highest levels achieved in this study.

Fc titers produced by KM_ultra_ were evaluated in a bioreactor. Supernatants were treated with PNGase F and quantified against β-lactoglobulin standards. With comparable biomass to WT, Fc secretion in KM_ultra_ increased continuously over time (**Figure 7H, I**) and remained consistently higher than WT. Fc reached 0.45 g/L in WT and 1.36 g/L in KM_ultra_ at 60 h, representing an ∼ 3-fold increase.

Given the substantial improvement in Fc production by KM_ultra_, we next quantified the production of three commercially successful Fc-fusion glycoproteins-etanercept, dulaglutide, and abatacept. Because the target proteins are heavily glycosylated and overlap with background bands in the supernatant, they were purified using Protein A beads and quantified by BCA assay. The identities of the purified proteins were confirmed by Western blotting, using an empty vector as a control (**Figure 8**). For dulaglutide, KM_ultra_ showed the most pronounced improvement, reaching a maximum purified concentration of 5.93 g/L at 60 h, compared with 0.49 g/L in WT (∼12-fold increase; **Figure 8A**). For abatacept, KM_ultra_ reached 7.57 g/L at 36 h, versus 2.14 g/L in WT at the same time point (∼3.5-fold increase; **Figure 8B**). Etanercept exhibited substantial degradation, likely at the Fc linkage. Nevertheless, the maximum purified concentration (including degraded forms) reached 5.32 g/L in KM_ultra_ and 4.09 g/L in WT at 48 h (∼1.3-fold increase, **Figure 8C**).

**Figure 8.**
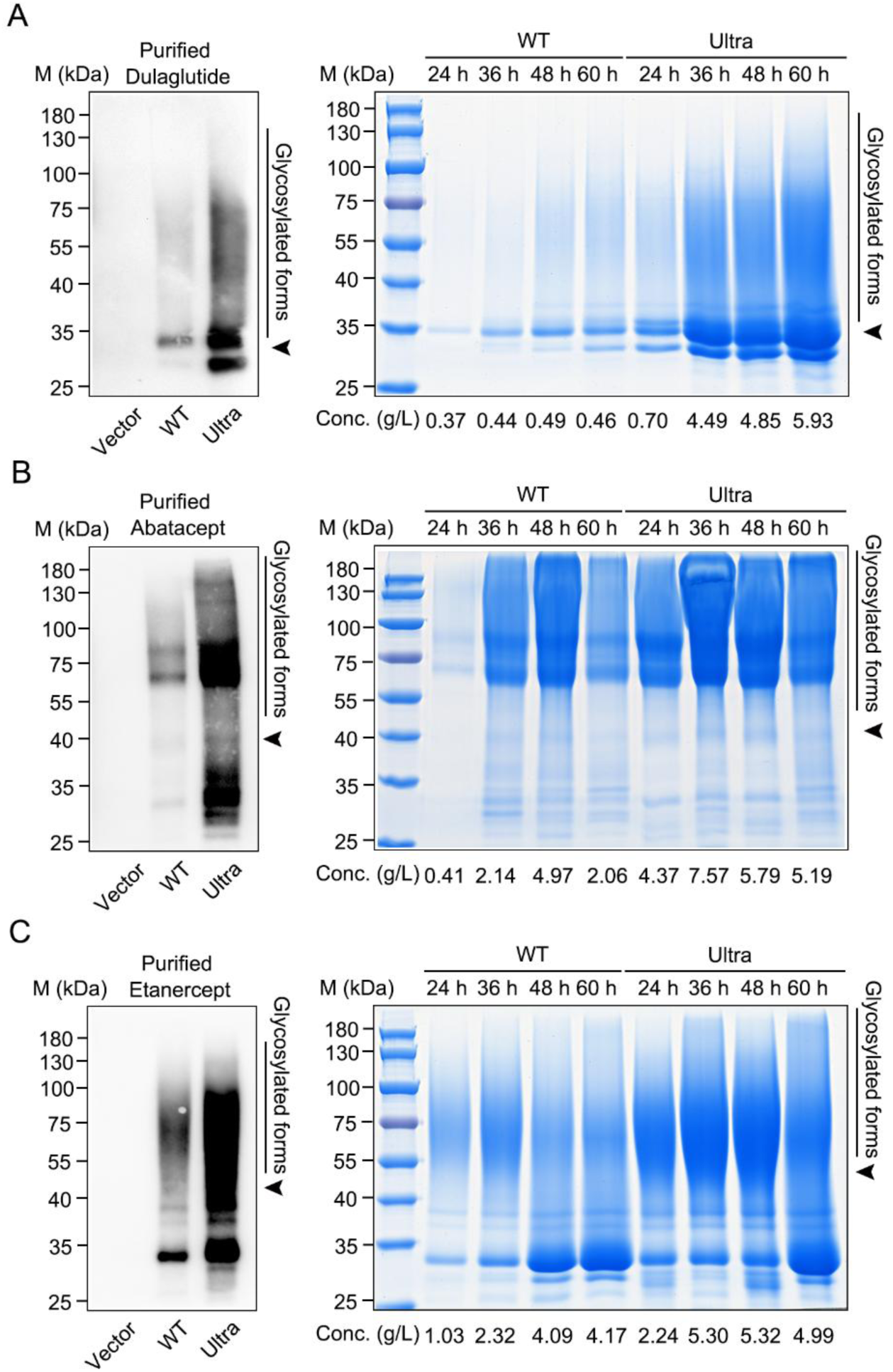
Humanized *K. marxianus* promotes high-level production of Fc-fusion therapeutics. (**A–C**) Production of dulaglutide (A), abatacept (B), and etanercept (C) in WT and KM_ultra_ (Ultra). Plasmids expressing dulaglutide (LHZ2070), abatacept (LHZ2071), or etanercept (LHZ2072) were introduced into WT and KM_ultra_. Supernatants were collected at the indicated times during bioreactor cultivation, purified (∼50 fold concentration), and analyzed by SDS–PAGE. Samples from the 24 h time point were diluted and analyzed by Western blotting using an anti-Fc antibody, with the 24 h sample from WT cells transformed with the empty vector (pUKDN132) as a control. Concentrations (Conc.) of purified proteins were determined by BCA assay and are shown below the corresponding lanes. Arrows indicate the positions of the target proteins at their expected molecular weights.

Together, these results demonstrate that the humanized KM_ultra_ strain has strong potential to enhance the production of diverse glycoprotein therapeutics.

## 3. Discussion

Compared with yeast, two key features of the human glycoprotein QC system, including iterative folding via the CNX/CRT cycle and stringent control of ERAD entry, prolong ER retention, and increase the probability of correct folding through repeated interactions with ER chaperones and folding enzymes. In this study, we introduced human UGGTs together with their cochaperone, as well as a human ERAD rate-limiting enzyme, into *K. marxianus*. In addition, we engineered a native GlsII trimming mutant. These modifications likely act by extending ER residence time, thereby rewiring the streamlined yeast QC system into a more humanized network and enhancing glycoprotein folding and secretion.

Heterologous expression of human UGGT1 or UGGT2 in *K. marxianus* enhanced glycoprotein secretion. Although we did not directly confirm UGGT activity by glycan profiling, the enhanced secretion likely depends on its enzymatic function. This is supported by three observations: (i) catalytic UGGT mutants failed to promote secretion (Figure 2E-I), (ii) co-expression of SEP15, which enhances UGGT activity, further increased secretion (Figure 3E-K), and (iii) UGGT-mediated secretion was abolished in the absence of GlsII activity (Figure 5D). Together, these results indicate that UGGT1/UGGT2 retain functional glucosyltransferase activity in *K. marxianus*. To date, the only heterologous UGGT shown to be enzymatically active in budding yeast is from *Schizosaccharomyces pombe* ^[33]^. Notably, human UGGT1 and UGGT2 share conserved functional domains with the S. pombe enzyme, with >60% identity in the UDP-glucose:glycoprotein glucosyltransferase domain (Figure S9). This suggests that, despite ∼1.3 billion years of divergence, UGGT proteins retain a robust catalytic core that functions across diverse systems.

In mammalian cells, G1M9 glycans, regenerated by UGGTs, are recognized by the CNX/CRT to facilitate protein folding. Consistently, UGGT1-mediated enhancement of substrate solubility depends on CRT and is abolished in its absence ^[34]^. However, our results suggest that UGGT can still improve folding efficiency in the absence of a canonical CNX/CRT system, as its effect is independent of Cne1, the yeast homolog of CRT (Figure 4C). *K. marxianus* retains functional GlsII activity (Figure 5C), and the coexistence of human UGGT and native GlsII may enable interconversion between G0M9 and G1M9, thereby prolonging ER retention. Even without CNX/CRT and their cofactors, endogenous ER chaperones such as Pdi1 and BiP/Kar2 may support folding ^[35, 36]^. Prolonged ER retention increases the probability of productive folding and enhances secretion. Consistently, introducing a native GlsII trimming mutant, which delays glucose removal and likely extends ER residence time ^[31]^, further enhanced UGGT1-mediated secretion (Figure 7C).

Introducing human EDEM2 into K. marxianus significantly delayed glycoprotein degradation in an activity-dependent manner (Figure 6C), indicating that EDEM2 retains its catalytic activity in this host. In mammalian cells, EDEM2 requires disulfide bonding with TXNDC11 to catalyze the conversion of G0M9 to M8B ^[37]^, whereas in yeast its homolog Htm1 forms a disulfide-linked complex with Pdi1 to generate a glycoprotein-specific mannosidase ^[38]^.

Although the overall sequence identity between TXNDC11 and Pdi1 is low (<10%), both proteins contain thioredoxin-fold domains, and EDEM2 retains conserved cysteines analogous to those mediating Htm1–Pdi1 disulfide bonding ^[37]^. These observations suggest that EDEM2 may similarly engage Pdi1 in yeast to catalyze the conversion of G0M9 to M8B, although this remains to be tested.

Mannose trimming by EDEM2 has been proposed to be slower than that by yeast Mns1^[15]^, raising the possibility that EDEM2 competes with Mns1 for G0M9 substrates and thereby delays M8B formation. Because downstream ERAD initiation by Htm1 depends on M8B^[39]^, EDEM2 expression likely attenuates ERAD initiation and subsequent degradation, thereby promoting protein secretion. Notably, EDEM2 enhanced protein secretion more effectively than *MNS1* deletion. While *MNS1* disruption can increase secretion in some cases^[40]^, it severely impairs ERAD, potentially leading to the accumulation of misfolded proteins that compromise folding capacity and secretion. In contrast, EDEM2 provides a moderated delay in ERAD, extending the folding window to improve secretion without incurring the detrimental effects of global ERAD disruption.

A limitation of this study is that we were unable to reconstitute a functional CRT/CNX cycle in *K. marxianus* by introducing human CRT/CNX and their cofactors. This was evidenced by the lack of improvement in glycoprotein expression across all tested combinations. Notably, co-expression of CRT/CNX with ERp57, ERp29, and CypB completely abolished the UGGT1-mediated enhancement of glycoprotein secretion, resulting in secretion levels even lower than those in WT (Figure 4K, L). These findings suggest that CRT/CNX and their cofactors fail to functionally cooperate with UGGT1 and may instead exert aberrant effects that impair glycoprotein folding and expression. Several factors could contribute to this dysfunction, including improper protein localization, abnormal substrate binding, or substrate trapping. One possible solution is to test CNX/CRT and their cofactors from species more closely related to *K. marxianus*, which may facilitate the reconstruction of a functional CRT/CNX cycle in yeast.

Finally, by combining five effective engineering strategies, we generated the KM_ultra_ strain. This strain further enhanced the production of Fc, Xyn-CDBFV, and GGH, with increases of up to 200%, and achieved a maximum Fc titer of 1.3 g/L. Notably, KM_ultra_ also improved the production of three high-value Fc-fusion therapeutics, with dulaglutide increasing by up to 12-fold. After accounting for concentration during purification, the original titers of Fc-fusion therapeutics in the supernatant exceeded 100 mg/L, which remains below those typically achieved in CHO cells (∼5 g/L). However, further improvements-such as glycoengineering to enhance product homogeneity^[5]^, deletion of vacuolar proteases to reduce degradation^[41]^, and optimization of fermentation conditions-are likely to increase titers to the gram-per-liter level. Given that *K. marxianus* has a short fermentation cycle (48–60 h), compared with 10–14 days for CHO fed-batch cultures, an optimized and humanized *K. marxianus* platform could provide a competitive alternative for producing high-value human glycoproteins.

## 4. Methods

### 4.1 Plasmids and primers

Plasmids and primers used in this study are listed in Tables S1 and S3, respectively. Donor plasmids (LHZ2030–LHZ2053) were constructed by ligating donor sequences into the pMD18-T vector (6011, Takara). Donors for gene insertion consisted of an upstream homologous arm (HA), an expression cassette (promoter-signal peptide-coding sequence [CDS]-HDEL–terminator), and a downstream HA, whereas donors for gene deletion contained only upstream and downstream HAs. In some constructs, specific components of the expression cassette were omitted depending on the target gene. Tandem expression cassettes were included in some constructs. The origins of donor plasmid components are described below.

The CDSs of *UGGT1*, *UGGT2*, *SEP15*, *EDEM2*, *CANX*, *CALR*, *PDIA3*, *ERP29*, *PPIB*, *GANAB*, and *PRKCSH*, with their native signal peptides removed, were amplified from HeLa cell cDNA. The *GTB1* CDS was amplified from the genome of FIM1ΔU. Point mutations in *UGGT1*, *UGGT2*, *EDEM2*, and *GTB1* were introduced by PCR-based mutagenesis. For heterologous expression, the sequence encoding the signal peptide of *K. marxianus* Pdi1 was fused to the N-terminus of human CDSs, except for CNX, and the ER-retention signal HDEL was fused to the C-terminus of each human CDS except CNX and UGGT2. The sequence encoding the Cne1 signal peptide was fused to the N-terminus of the CNX CDS. Promoters and terminators of *S. cerevisiae* (*ADH1*, *ADH2*, *PGK1*, *GPD1*, *TDH3*, and *TEF1*) were amplified from the S288C genome, whereas the *AFT1* promoter and terminator from *P. pastoris* were amplified from the GS115 genome. Upstream and downstream sequences of the Sg1, Sg4, Sg7, Sg19, *GTB1*, *ROT2*, *MNS1* and *CNE1* loci were used as homologous arms. For sequential genomic integrations, 5’ regions of the *ScADH2*, *ScTEF2*, and *ScPGK1* promoters were used as downstream HAs to target insertion upstream of previously integrated expression cassettes. Detailed compositions of donor plasmids, including Gene IDs, HA lengths, promoters, signal peptides, and terminators, are provided in Table S1.

For CRISPR genome editing, a Sta2-Cas9 empty vector (LHZ2065) was constructed by replacing the SpCas9 ORF in LHZ531 ^[42]^ with Sta2-Cas9, and substituting the original gRNA sequence between the tRNA-Gly element and the SUP4 terminator with a filler fragment containing two AarI restriction sites and a gRNA scaffold compatible with the Sta2-Cas9 system. The filler and scaffold sequences are listed in Table S3. Target sequences (21 bp) were introduced by annealing primer pairs and inserting them into the AarI sites of LHZ2065 to generate plasmids LHZ2066 and LHZ2067. For SpCas9 constructs (LHZ2054–LHZ2064), primer pairs containing 20-bp target sequences were inserted into the SapI sites of the corresponding vector.

For secretory expression, the CDSs of Fc or GGH with a C-terminal His₆ tag, and the CDSs of dulaglutide, abatacept, and etanercept (https://go.drugbank.com), were inserted between the SmaI and NotI sites of LHZ399 (pZP33) ^[27]^, generating plasmids LHZ2068–LHZ2072, respectively. The empty vector (pUKDN132) and LHZ443 (pZP46) overexpressing Xyn-CDBFV was described previously ^[27]^. The sequences encoding dulaglutide, abatacept, and etanercept were obtained from DrugBank (https://go.drugbank.com), codon-optimized, synthesized, and inserted between the SmaI and NotI sites of LHZ399, generating the plasmids LHZ2070, LHZ2071, and LHZ2072, respectively.

### 4.2 Strains and media

Strains used in this study are listed in Table S2. Strains were cultivated at 30 °C in YPD medium (2% peptone, 1% yeast extract, 2% glucose; 2% agar for plates) for routine culturing, in synthetic complete medium lacking uracil (SC-Ura) for selecting transformants, or in YD medium (2% yeast extract, 4% glucose) for secretory expression.

FIM1ΔU^[27]^ was used as the wild-type strain. Strains were constructed by transforming a CRISPR plasmid and a donor fragment amplified from a donor plasmid into the host strain. Transformants were selected on SC-Ura plates. Details of the CRISPR plasmid, donor plasmid, and host strain used for each constructed strain are listed in Table S2. In brief, *UGGT1*, *UGGT1-D1358A*, *UGGT2*, *UGGT2-D1333A*, *SEP15*, *EDEM2*, *EDEM2-E117Q*, *CANX*, *CALR*, *ERP57*, and *PPIB*-*ERP29* were introduced into FIM1ΔU to generate KM_UGGT1_, KM_UGGT1-D1358A_, KM_UGGT2_, KM_U2-D1333A_, KM_SEP15_, KM_EDEM2_, KM_E-E117Q_, KM_CNX_, KM_CRT_,

KM_p57_, and KM_cypb-p29_, respectively. *UGGT2* was introduced into KM_UGGT1_ to generate KM_U1&2_. *SEP15* was introduced into KM_UGGT1_, KM_UGGT2_, and KM_U1&2_ to generate KM_U1-S_, KM_U2-S_, and KM_U1&2-S_, respectively. *ERP57* was introduced into KM_CNX_ and KM_CRT_ to generate KM_CNX-p57_ and KM_CRT-p57_, respectively. *PPIB*-*ERP29* was introduced into KM_CNX-p57_ and KM_CRT-p57_ to generate KM_CNX-3C_ and KM_CRT-3C_, respectively. *GTB1* was deleted in FIM1ΔU and KM_U1&2-S_ to generate KM_gtb1Δ_ and KM_U1&2-S-gtb1Δ_, respectively. *GTB1*-*E110A*, *GTB1*-*R629A*, and *GTB1*-*Y654F* were introduced into KM_gtb1Δ_ to generate KM_β-E110A_, KM_β-R629A_, and KM_β-Y654F_, respectively. *GTB1*-*E110A* was introduced into KM_U1&2-S-gtb1Δ_ to generate KM_U1&2-S-E110A_. *CNE1* was deleted in FIM1ΔU and KM_UGGT1_ to generate KM_cne1Δ_ and KM_U1-cne1Δ_, respectively. *ROT2* was deleted in FIM1ΔU and KM_gtb1Δ_ to generate KM_rot2Δ_ and KM_GIIΔ_, respectively. *GANAB* was introduced into KM*_rot2_*_Δ_ to generate KM_hGIIα_, and *PRKCSH* was introduced into KM_hGIIα_ to generate KM_hGII_. *MNS1* was deleted in FIM1ΔU to generate KM_mns1Δ_. *EDEM2* was introduced into KM_U1&2-S_ and KM_U1&2-S-E110A_ to generate KM_U1&2-S-EDEM2_ and KM_ultra_, respectively.

### 4.3 Detection of human glycoprotein quality-control proteins

Yeast cells were inoculated into 50 mL YPD medium at an initial OD_600_ of 0.2 and cultured for 4–6 h at 30 °C until the exponential phase. Cells equivalent to 75 OD_600_ units were harvested and resuspended in 5 mL PBS buffer (pH 7.2). One protease inhibitor cocktail tablet (Roche, 04693159001) was dissolved in 10 mL ddH₂O to prepare the protease inhibitor working solution. The PBS-resuspended cells were centrifuged again, and the resulting cell pellet was resuspended in 500 μL of the protease inhibitor working solution. The cell suspension was mixed with 500 μL glass beads (G8772, Sigma-Aldrich) and disrupted using a bead beater (FastPrep-24, MP) at 6 m/s for a total of 3 min, with cooling on ice for 2 min after every 30 s of processing. The lysate was centrifuged at 12,000 rpm for 20 min at 4 °C, and the supernatant was collected. The supernatant or pellet was analyzed by SDS-PAGE or by native PAGE (Molecular Cloning). For native PAGE, SDS was omitted from the SDS-PAGE reagents. For Western blot (WB) analysis, proteins were transferred onto PVDF membranes after electrophoresis and detected using primary antibodies against UGGT1 (14170-1-AP), UGGT2 (13420-1-AP), EDEM2 (11241-1-AP), CNX (10427-2-AP), CRT (10292-1-AP), ERp57 (15967-1-AP), ERp29 (24344-1-AP), CypB (11607-1-AP), GANAB (83324-1-RR), PRKCSH (12148-1-AP), Fc (68275-1-Ig), and β-actin (20536-1-AP) (all from Proteintech), as well as SEP15 (R25690, ZENBIO) and GAPDH (PT0582R, Immunoway). HRP-conjugated secondary antibodies (5220-0341 and 5220-0336, SeraCare) were used, and signals were visualized using an enhanced chemiluminescence (ECL) detection reagent (RPN2232, Cytiva).

### 4.4 Growth curve analysis

Overnight cultures were inoculated into 50 mL YPD medium at an initial OD₆₀₀ of 0.1 and cultured for 24 h. OD₆₀₀ values were measured at the indicated time points. Experiments were performed with three parallel cultures.

### 4.5 Expression of glycoproteins

Cells were transformed with LHZ2068 (Fc), LHZ2069 (GGH), or LHZ443 (Xyn), and transformants were selected on SC-Ura plates. For GGH and Xyn expression, transformants were cultured in 50 mL YD medium for 72 h. Culture supernatants from Xyn samples were analyzed by SDS-PAGE, whereas those from GGH samples were analyzed by SDS-PAGE followed by WB using an anti-His₆ antibody (66005-1-Ig, Proteintech).

For intracellular Fc detection, transformants were cultured in 50 mL YD medium for 72 h. Cells equivalent to 15 OD_600_ units were harvested and lysed using the glass bead method described above. After centrifugation, the supernatant was collected as the soluble fraction. The pellet was resuspended in 1×SDS loading buffer to the same volume as the lysate, boiled, and centrifuged, and the supernatant was collected as the insoluble fraction. The soluble fraction and a 2,000-fold diluted insoluble fraction were analyzed by SDS-PAGE followed by WB using an anti-His₆ antibody.

For large-scale production of Fc, dulaglutide, abatacept, and etanercept, transformants were subjected to fed-batch fermentation. A seed culture was prepared by growing cells in 50 mL SM medium (0.5% (NH₄)₂SO₄, 1% glucose, 0.05% MgSO₄·7H₂O, 0.3% KH₂PO₄) in a 250 mL flask at 30 °C and 220 rpm for 16 h. The seed culture was then inoculated into a 1.5 L Minibox Intelli-Ferm 6 bioreactor (T&J Bio-engineering) containing 600 mL SM medium.

Fermentation was carried out at 30 °C with sterile air supplied at 0.6 L/min. Dissolved oxygen (DO) was maintained above 10% by adjusting the agitation speed between 300 and 1000 rpm. The pH was maintained approximately 5.5 using ammonia. Glucose was continuously fed as a 650 g/L solution, with a total of 300 g supplied during fermentation. Culture supernatants were collected at 24, 36, 48, 60, and 72 h. Cell pellets were dried at 80 °C for 24 h to determine dry cell weight.

### 4.6 Quantification of protein levels and mobility shift

SDS-PAGE gels were stained with Coomassie Brilliant Blue G-250 and scanned using an Epson Perfection V600 Photo scanner. WB membranes were imaged using a GeneGnome Bio Imaging System (Syngene). Band intensities were quantified using ImageJ to estimate relative protein levels among samples.

For fed-batch fermentation samples of Fc, supernatants were analyzed by 15% (w/v) SDS-PAGE together with serial dilutions of β-lactoglobulin (L3908, Sigma-Aldrich). Gels were scanned and β-lactoglobulin band intensities were used to generate a standard curve correlating band intensity with protein concentration. The concentrations of target proteins were then calculated from this standard curve.

For fed-batch fermentation samples of dulaglutide, abatacept, and etanercept, proteins were purified from clarified supernatants using rProtein A beads (SA012025, Smart-Lifesciences) according to the manufacturer’s protocol. A total of 600 μL of eluate was obtained from 30 mL of supernatant (∼50-fold concentration). Protein concentrations were determined using a BCA assay kit (C503021-0500, Sangon).

To quantify band mobility, grayscale intensities along each lane (bottom to top) were measured using ImageJ and binned into small intervals. The data were then smoothed and visualized as ridge plots in R (version 4.3.1) using ggplot2. The median migration position (indicated by a vertical line) was defined as the point where cumulative signal reached 50% of the total within the analyzed lane.

### 4.7 PNGase F deglycosylation analysis

Supernatants were subjected to deglycosylation using PNGase F (20407ES01, Yeasen) according to the manufacturer’s instructions. Reactions were incubated at 37 °C for 1 h. After termination, samples were analyzed by SDS–PAGE (for Fc and Xyn) or by WB (for γ-glutamyl hydrolase).

### 4.8 Cycloheximide (CHX) chase degradation assay

Overnight cultures were inoculated into 50 mL YPD medium at an initial OD₆₀₀ of 0.1 and cultured for 24 h. Cells were harvested and resuspended in pre-warmed fresh YD medium. Samples collected immediately were designated as the 0 min time point. CHX (MCE, HY-12320) was then added to a final concentration of 250 μg/mL to inhibit protein synthesis. Samples were collected at 10, 15, 30, 60, 120, and 180 min after CHX addition. At each time point, cells equivalent to 15 OD₆₀₀ units were harvested, washed once with ice-cold PBS, and lysed using the glass bead method as described above. Supernatants were analyzed by SDS–PAGE followed by WB using an anti-His antibody. Band intensities were quantified using ImageJ. Protein signals at each time point were normalized to the signal at 0 min (set as 100%). Experiments were performed with three biological replicates.

## Supporting information

Supplementary Table 1. Plasmids used in this study

Supplementary Table 2. Strains used in this study

Supplementary Table 3. Primers and synthetic DNA sequences used in this study

## Acknowledgements

This work was supported by National Key Research and Development Program of China (2021YFA0910601, 2021YFA0910603), and Science and Technology Research Program of Shanghai (24HC2810100, 24ZR1406500, 2023ZX01).

## Data Availability Statement

The data supporting the findings of this study are available within the article and its Supplementary Information.

## References

[1] Chung C Y, Majewska N I, Wang Q, et al. SnapShot: N-Glycosylation Processing Pathways across Kingdoms [J]. Cell, 2017, 171(1): 258–258.e251.

[2] O’flaherty R, Bergin A, Flampouri E, et al. Mammalian cell culture for production of recombinant proteins: A review of the critical steps in their biomanufacturing [J]. Biotechnol Adv, 2020, 43: 107552.

[3] Baghban R, Farajnia S, Rajabibazl M, et al. Yeast Expression Systems: Overview and Recent Advances [J]. Mol Biotechnol, 2019, 61(5): 365–384.

[4] Wildt S, Gerngross T U. The humanization of N-glycosylation pathways in yeast [J]. Nat Rev Microbiol, 2005, 3(2): 119–128.

[5] Piirainen M A, Frey A D. The Impact of Glycoengineering on the Endoplasmic Reticulum Quality Control System in Yeasts [J]. Front Mol Biosci, 2022, 9: 910709.

[6] Ye J, Ly J, Watts K, et al. Optimization of a glycoengineered Pichia pastoris cultivation process for commercial antibody production [J]. Biotechnol Prog, 2011, 27(6): 1744–1750.

[7] Lin H, Kim T, Xiong F, et al. Enhancing the production of Fc fusion protein in fed-batch fermentation of Pichia pastoris by design of experiments [J]. Biotechnol Prog, 2007, 23(3): 621–625.

[8] Huang Y M, Hu W, Rustandi E, et al. Maximizing productivity of Cho cell-based fed-batch culture using chemically defined media conditions and typical manufacturing equipment [J]. Biotechnol Prog, 2010, 26(5): 1400–1410.

[9] Ying B, Kawabe Y, Zheng F, et al. High-Level Production of scFv-Fc Antibody Using an Artificial Promoter System with Transcriptional Positive Feedback Loop of Transactivator in Cho Cells [J]. Cells, 2023, 12(22).

[10] Guay K P, Chou W C, Canniff N P, et al. N-glycan-dependent protein maturation and quality control in the ER [J]. Nat Rev Mol Cell Biol, 2025, 26(12): 926–939.

[11] Kozlov G, Gehring K. Calnexin cycle - structural features of the ER chaperone system [J]. Febs j, 2020, 287(20): 4322–4340.

[12] Williams R V, Guay K P, Hurlbut Lesk O A, et al. Insights into the interaction between UGGT, the gatekeeper of folding in the ER, and its partner, the selenoprotein SEP15 [J]. Proc Natl Acad Sci U S A, 2024, 121(34): e2315009121.

[13] Parodi A, Cummings R D, Aebi M. Glycans in Glycoprotein Quality Control [M]//Varki A, Cummings R D, Esko J D, et al. Essentials of Glycobiology. Cold Spring Harbor (NY); Cold Spring Harbor Laboratory Press Copyright 2015-2017 by The Consortium of Glycobiology Editors, La Jolla, California. All rights reserved. 2015: 503-511.

[14] Banerjee S, Vishwanath P, Cui J, et al. The evolution of N-glycan-dependent endoplasmic reticulum quality control factors for glycoprotein folding and degradation [J]. Proc Natl Acad Sci U S A, 2007, 104(28): 11676–11681.

[15] Ninagawa S, Okada T, Sumitomo Y, et al. EDEM2 initiates mammalian glycoprotein ERAD by catalyzing the first mannose trimming step [J]. J Cell Biol, 2014, 206(3): 347–356.

[16] Pfeffer M, Maurer M, Stadlmann J, et al. Intracellular interactome of secreted antibody Fab fragment in Pichia pastoris reveals its routes of secretion and degradation [J]. Appl Microbiol Biotechnol, 2012, 93(6): 2503–2512.

[17] Ciplys E, Sasnauskas K, Slibinskas R. Overexpression of human calnexin in yeast improves measles surface glycoprotein solubility [J]. Fems Yeast Res, 2011, 11(6): 514–523.

[18] Duan J, Yang D, Chen L, et al. Efficient production of porcine circovirus virus-like particles using the nonconventional yeast Kluyveromyces marxianus [J]. Appl Microbiol Biotechnol, 2019, 103(2): 833–842.

[19] Yang D, Chen L, Duan J, et al. Investigation of Kluyveromyces marxianus as a novel host for large-scale production of porcine parvovirus virus-like particles [J]. Microb Cell Fact, 2021, 20(1): 24.

[20] Tian T, Wu X, Wu P, et al. High-level expression of leghemoglobin in Kluyveromyces marxianus by remodeling the heme metabolism pathway [J]. Front Bioeng Biotechnol, 2023, 11: 1329016.

[21] Wu P, Mo W, Tian T, et al. Transfer of disulfide bond formation modules via yeast artificial chromosomes promotes the expression of heterologous proteins in Kluyveromyces marxianus [J]. mLife, 2024, 3(1): 129–142.

[22] Lee M H, Hsu T L, Lin J J, et al. Constructing a human complex type N-linked glycosylation pathway in Kluyveromyces marxianus [J]. PLos One, 2020, 15(5): e0233492.

[23] Karbalaei M, Rezaee S A, Farsiani H. Pichia pastoris: A highly successful expression system for optimal synthesis of heterologous proteins [J]. J Cell Physiol, 2020, 235(9): 5867–5881.

[24] Assunção Bicca S, Poncet-Legrand C, Williams P, et al. Structural characteristics of Saccharomyces cerevisiae mannoproteins: Impact of their polysaccharide part [J]. Carbohydr Polym, 2022, 277: 118758.

[25] Takeda Y, Seko A, Hachisu M, et al. Both isoforms of human UDP-glucose:glycoprotein glucosyltransferase are enzymatically active [J]. Glycobiology, 2014, 24(4): 344–350.

[26] Adams B M, Canniff N P, Guay K P, et al. Quantitative glycoproteomics reveals cellular substrate selectivity of the ER protein quality control sensors UGGT1 and UGGT2 [J]. Elife, 2020, 9.

[27] Zhou J, Zhu P, Hu X, et al. Improved secretory expression of lignocellulolytic enzymes in Kluyveromyces marxianus by promoter and signal sequence engineering [J]. Biotechnol Biofuels, 2018, 11: 235.

[28] Ren B, Liu M, Ni J, et al. Role of Selenoprotein F in Protein Folding and Secretion: Potential Involvement in Human Disease [J]. Nutrients, 2018, 10(11).

[29] Kurita T, Noda Y, Yoda K. Action of multiple endoplasmic reticulum chaperon-like proteins is required for proper folding and polarized localization of Kre6 protein essential in yeast cell wall β-1,6-glucan synthesis [J]. J Biol Chem, 2012, 287(21): 17415–17424.

[30] Wilkinson B M, Purswani J, Stirling C J. Yeast GTB1 encodes a subunit of glucosidase Ii required for glycoprotein processing in the endoplasmic reticulum [J]. J Biol Chem, 2006, 281(10): 6325–6333.

[31] Quinn R P, Mahoney S J, Wilkinson B M, et al. A novel role for Gtb1p in glucose trimming of N-linked glycans [J]. Glycobiology, 2009, 19(12): 1408–1416.

[32] Hosomi A, Tanabe K, Hirayama H, et al. Identification of an Htm1 (EDEM)-dependent, Mns1-independent Endoplasmic Reticulum-associated Degradation (ERAD) pathway in Saccharomyces cerevisiae: application of a novel assay for glycoprotein Erad [J]. J Biol Chem, 2010, 285(32): 24324–24334.

[33] Castro O, Chen L Y, Parodi A J, et al. Uridine diphosphate-glucose transport into the endoplasmic reticulum of Saccharomyces cerevisiae: in vivo and in vitro evidence [J]. Mol Biol Cell, 1999, 10(4): 1019–1030.

[34] Ferris S P, Jaber N S, Molinari M, et al. UDP-glucose:glycoprotein glucosyltransferase (UGGT1) promotes substrate solubility in the endoplasmic reticulum [J]. Mol Biol Cell, 2013, 24(17): 2597–2608.

[35] Frand A R, Kaiser C A. Ero1p oxidizes protein disulfide isomerase in a pathway for disulfide bond formation in the endoplasmic reticulum [J]. Mol Cell, 1999, 4(4): 469–477.

[36] Simons J F, Ferro-Novick S, Rose M D, et al. BiP/Kar2p serves as a molecular chaperone during carboxypeptidase Y folding in yeast [J]. J Cell Biol, 1995, 130(1): 41–49.

[37] George G, Ninagawa S, Yagi H, et al. EDEM2 stably disulfide-bonded to TXNDC11 catalyzes the first mannose trimming step in mammalian glycoprotein Erad [J]. Elife, 2020, 9.

[38] Liu Y C, Fujimori D G, Weissman J S. Htm1p-Pdi1p is a folding-sensitive mannosidase that marks N-glycoproteins for ER-associated protein degradation [J]. Proc Natl Acad Sci U S A, 2016, 113(28): E4015–4024.

[39] Clerc S, Hirsch C, Oggier D M, et al. Htm1 protein generates the N-glycan signal for glycoprotein degradation in the endoplasmic reticulum [J]. J Cell Biol, 2009, 184(1): 159–172.

[40] Wu X, Feng D, Chen H, et al. Multistep engineering of the secretory pathway for enhanced leghemoglobin expression in Kluyveromyces marxianus [J]. Bioresour Technol, 2026, 448: 134275.

[41] Wu M, Shen Q, Yang Y, et al. Disruption of YPS1 and PEP4 genes reduces proteolytic degradation of secreted HSA/PTH in Pichia pastoris GS115 [J]. J Ind Microbiol Biotechnol, 2013, 40(6): 589–599.

[42] Zhou H, Tian T, Liu J, et al. Efficient and markerless gene integration with SlugCas9-HF in Kluyveromyces marxianus [J]. Commun Biol, 2024, 7(1): 797.

