## Supplemental Figures for "Humanization of N-glycan-dependent protein quality control system in *Kluyveromyces marxianus* promotes glycoprotein secretion"

**Figure S1. Verification of Fc and GGH glycosylation**

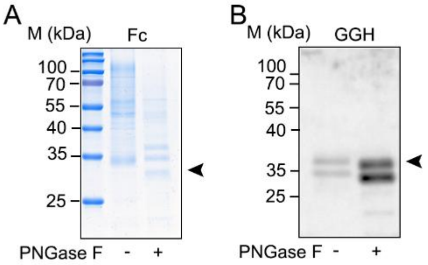


(A) Fc glycosylation. The Fc plasmid (LHZ2068) was transformed into WT. The transformant was cultured in a bioreactor for 24 h. Supernatants, with or without PNGase F treatment, were analyzed by SDS–PAGE.

(B) GGH glycosylation. The GGH plasmid (LHZ2069) was transformed into WT. The transformant was cultured in shake flasks for 72 h. Supernatants, with or without PNGase F treatment, were analyzed by WB using an anti-$His_6$ antibody.
Arrows indicate the positions of the target proteins at their expected molecular weights.

**Figure S2. Validation of band specificity for Fc, GGH, and Xyn-CDBFV**

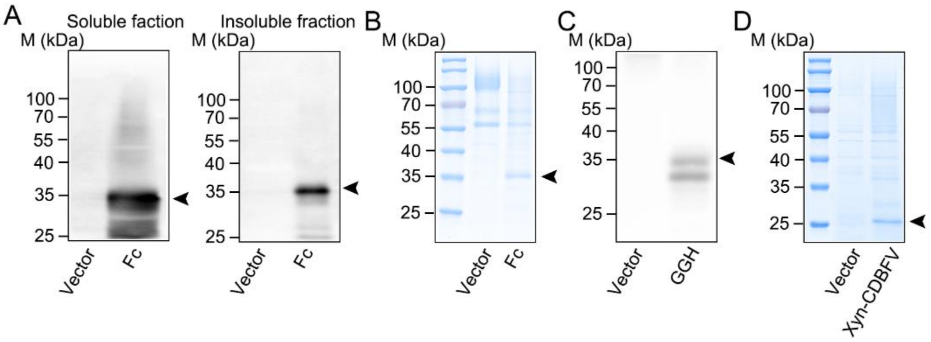


(A) Fc WB specificity. WT cells carrying the Fc plasmid (LHZ2068) or empty vector (pUKDN132) were cultured for 72 h. Cells were lysed and both supernatant and pellet (2000-fold dilution) fractions were analyzed by WB (anti-$His_6$). No non-specific bands were detected in vector controls. Black arrows here and below indicate the target proteins.

(B) Fc SDS-PAGE specificity. WT cells carrying LHZ2068 or empty vector were cultured for 24 h in a bioreactor. Supernatants were analyzed by SDS-PAGE. No bands were detected at the Fc position in controls.

(C) GGH WB specificity. WT cells carrying the GGH plasmid (LHZ2069) or empty vector were cultured for 72 h. Supernatants were analyzed by WB (anti-$His_6$). No non-specific bands were detected in controls.

(D) Xyn-CDBFV SDS-PAGE specificity. WT cells carrying the Xyn-CDBFV plasmid LHZ443 or empty vector were cultured for 72 h. Supernatants were analyzed by SDS-PAGE. No bands were detected at the Xyn-CDBFV position in controls.

**Figure S3. Effects of UGGT1 on glycoprotein secretion after PNGase F treatment**

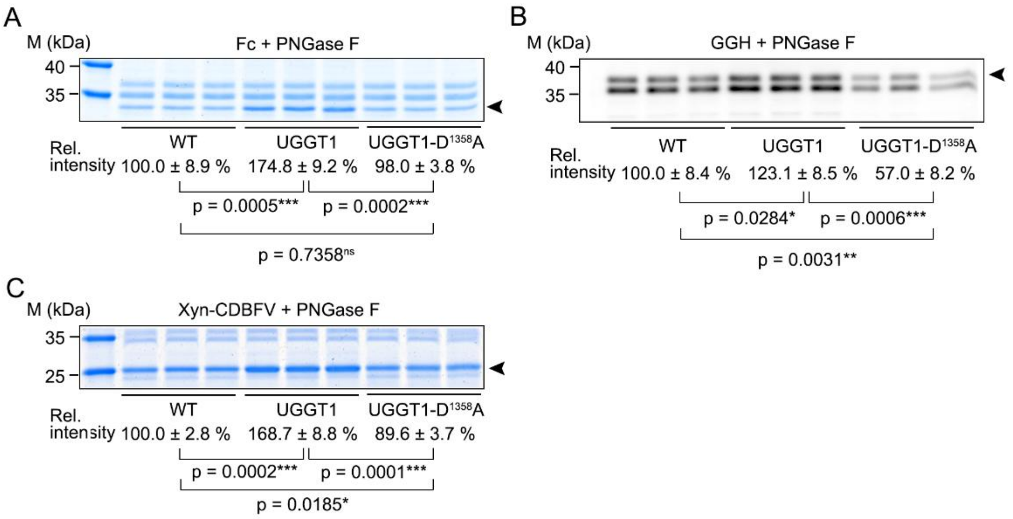


(A) Fc secretion. The Fc plasmid (LHZ2068) was introduced into WT, $KM_{UGGT1}$, or $KM_{U1\text{-}D1358A}$. After 24 h of bioreactor cultivation, supernatants were treated with PNGase F and analyzed by SDS-PAGE.

(B) GGH secretion. The GGH plasmid (LHZ2069) was introduced, and supernatants were analyzed after PNGase F treatment by WB (anti-$His_6$).

(C) Xyn-CDBFV secretion. The Xyn-CDBFV plasmid (LHZ443) was introduced, and supernatants were analyzed after PNGase F treatment by SDS-PAGE.

Arrows indicate the position of the target protein at its expected molecular weight. Band intensities were quantified using ImageJ, with WT set to 100%. Data are mean ± SD (n = 3).

**Figure S4. Effects of GlsII mutants on Fc mobility shift and secretion (replicate).**

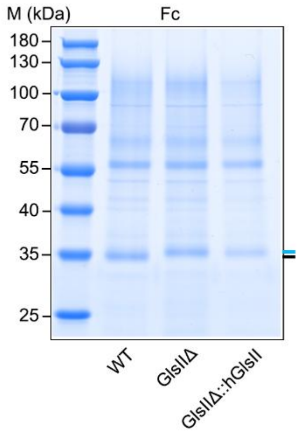


The Fc plasmid (LHZ2068) was introduced into WT, $KM_{GII\Delta}$ (GlsIIΔ), and $KM_{hGII}$ (GlsIIΔ::hGlsII). After 24 h of bioreactor cultivation, supernatants were analyzed by SDS-PAGE. Blue and black lines indicate Fc bands with or without mobility shift, respectively.

**Figure S5. Comparison of conserved amino acids in Gtb1 between *S. cerevisiae* and K. *marxianus*.**

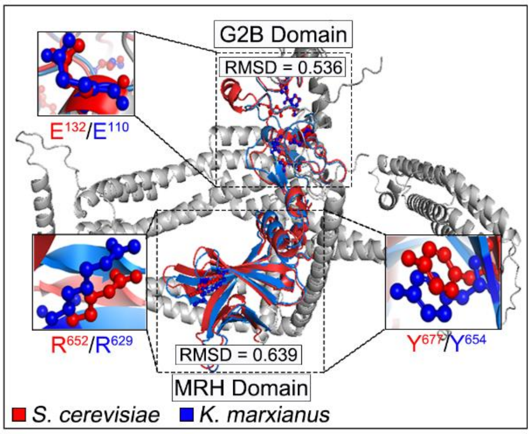


The structures of Gtb1 from *S. cerevisiae* and *K. marxianus* were predicted using AlphaFold. Structural superposition was performed in PyMOL using the align command, and the root mean square deviation (RMSD) is shown. Conserved amino acids essential for the catalytic activity of Gtb1 in *S. cerevisiae* (red) were used to identify the corresponding residues in *K. marxianus* (blue).

**Figure S6. Effect of EDEM2 on Fc degradation (replicates).**

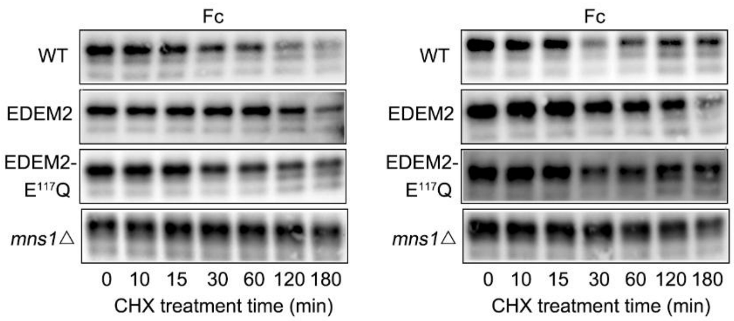


The Fc plasmid (LHZ2068) was introduced into WT, $KM_{EDEM2}$, $KM_{E\text{-}E117Q}$, and $KM_{mns1\Delta}$. After 24 h of shake-flask cultivation, cells were treated with cycloheximide (CHX) for the indicated times. Cell lysates were analyzed by WB using an anti-$His_6$ antibody.

**Figure S7. Fc titer in WT and $KM_{ultra}$ (without PNGase F treatment)**

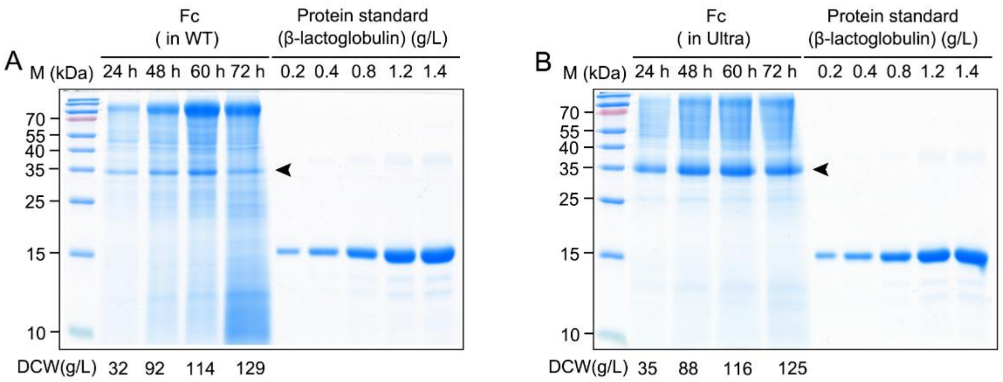


The Fc plasmid (LHZ2068) was introduced into WT (A) and $KM_{ultra}$ (B). Supernatants were collected after 24, 48, 60, and 72 h of bioreactor cultivation and analyzed by SDS-PAGE without PNGase F treatment.

**Figure S8. Standard curves of β-lactoglobulin.**

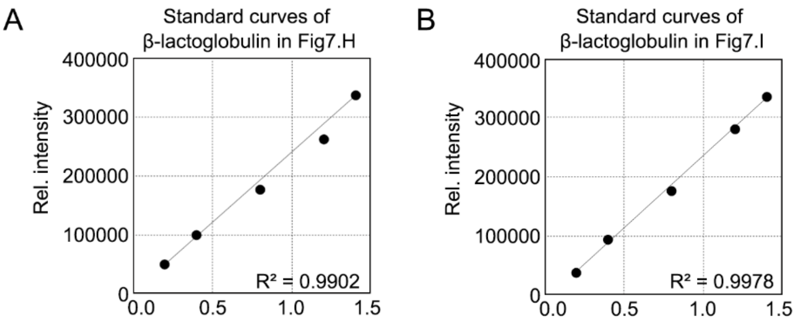


Standard curves were generated by grayscale analysis of β-lactoglobulin for quantification of Fc in Fig. 7H (A) and I (B). The coefficient of determination ($R^2$) is shown.

**Figure S9. Sequence identities between conserved domains of human and *S. pombe* UGGTs.**

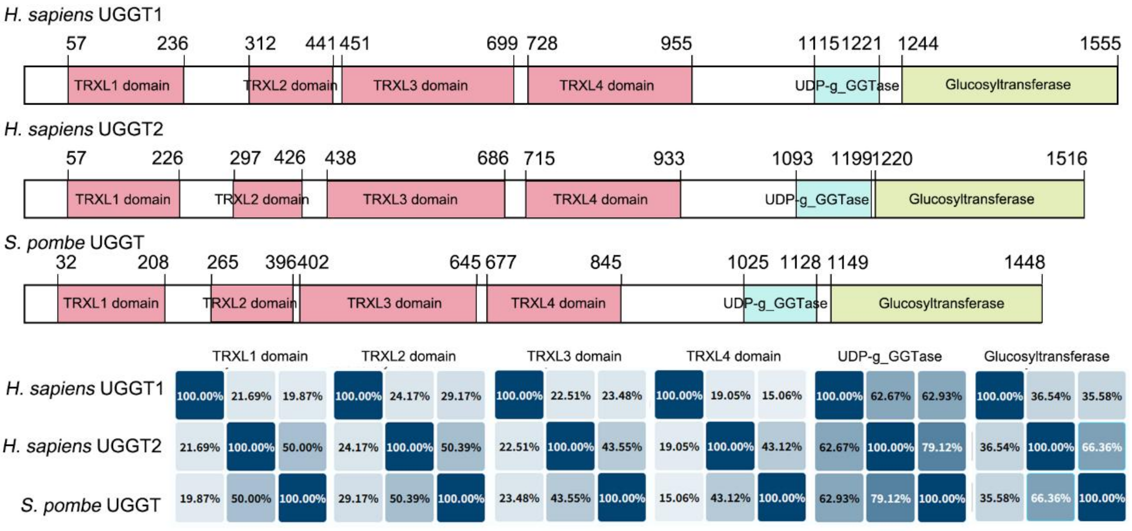


Domains were predicted using UniProt. The identity matrix (percentages) for each domain is shown below the alignment. UDP-g_GGTase: UDP-glucose:glycoprotein glucosyltransferase.
