## Supplementary Table 1. Plasmids used in this study for "Humanization of N-glycan-dependent protein quality control system in *Kluyveromyces marxianus* promotes glycoprotein secretion"

Table S1. Plasmids used in this study.

| Name | Essential Features | Backbone | Description and application | Source or reference |
|---|---|---|---|---|
| LHZ2030 | *AmpR*, Sg1 upstream-$P_{ScADH1}$-$SP_{KmPDI1}$-*HsUGGT1-HDEL*-$T_{ScADH1}$-Sg1 downstream | pMD-18T | 500 bp Sg1 upstream sequence, *ScADH1*[1] promoter[2] (Gene ID 854068), *KmPDI1* signal sequence (Gene ID 34713733, 1-63 bp), CDS of *HsUGGT1* (Gene ID 56886, 129-4653 bp), 1000 bp *ScADH1* terminator[3], 500 bp Sg1 downstream sequence were linked together. The resulting fragment was ligated into the pMD-18T vector to generate a donor plasmid for insertion of the *UGGT1* cassette into the Sg1 locus of FIM1ΔU. | this study |
| LHZ2031 | *AmpR*, Sg1 upstream-$P_{ScADH1}$-$SP_{KmPDI1}$-*HsUGGT1-D1358A-HDEL*-$T_{ScADH1}$-Sg1 downstream | pMD-18T | All elements were identical to those in LHZ2030, except that the *HsUGGT1* CDS carried the D1358A substitution. The resulting fragment was ligated into the pMD-18T vector to generate a donor plasmid for insertion of the *UGGT1-D1358A* cassette into the Sg1 locus of FIM1ΔU. | this study |
| LHZ2032 | *AmpR*, Sg4 upstream-$P_{ScGPD1}$-$SP_{KmPDI1}$-*HsUGGT2*-$T_{ScGPD1}$-Sg4 downstream | pMD-18T | 500 bp Sg4 upstream sequence, *ScGPD1*[1] promoter[2] (Gene ID 90072929), *KmPDI1* signal sequence, CDS of *HsUGGT2* (Gene ID 55757, 84-4548 bp), 1000 bp *ScGPD1* terminator[3], 500 bp Sg4 downstream sequence were linked together. The resulting fragment was ligated into the pMD-18T vector to generate a donor plasmid for | this study |

| | | | | |
|---|---|---|---|---|
| | | | insertion of the *UGGT2* cassette into the Sg4 locus of FIM1ΔU. | |
| LHZ2033 | *AmpR*, Sg4 upstream-P$_{ScGPD1}$-SP$_{KmPDI1}$-*HsUGGT2-D1333A*-T$_{ScGPD1}$-Sg4 downstream | pMD-18T | All elements were identical to those in LHZ2032, except that the *HsUGGT2* CDS carried the D1333A substitution. The resulting fragment was ligated into the pMD-18T vector to generate a donor plasmid for insertion of the *UGGT2-D1333A* cassette into the Sg4 locus of FIM1ΔU. | this study |
| LHZ2034 | *AmpR*, Sg7 upstream-P$_{ScCYS3}$ -SP$_{KmPDI1}$-*HsSEP15-U96C-HDEL*-T$_{ScCYS3}$-Sg7 downstream | pMD-18T | 500 bp Sg7 upstream sequence, *ScCYS3*[1] promoter[2] (Gene ID 851221), *KmPDI1* signal sequence, CDS of *HsSEP15-U96C* derived from SEP15 (Gene ID: 9403, 96–495 bp), 1000 bp *ScCYS3* terminator[3], 500 bp Sg7 downstream sequence were linked together. The resulting fragment was ligated into the pMD-18T vector to generate a donor plasmid for insertion of the *SEP15-U96C* cassette into the Sg7 locus of FIM1ΔU, KM$_{UGGT1}$, KM$_{UGGT2}$ and KM$_{U1\&2}$. | this study |
| LHZ2035 | *AmpR*, Sg19 upstream-P$_{ScTEF1}$-SP$_{KmPDI1}$-*HsEDEM2-HDEL*-T$_{ScTEF1}$-Sg19 downstream | pMD-18T | 500 bp Sg19 upstream sequence, *ScTEF1*[1] promoter[2] (Gene ID 856195), *KmPDI1* signal sequence, CDS of *HsEDEM2* (Gene ID 55741, 66-1734 bp), 1000 bp *ScTEF1* terminator[3], 500 bp Sg19 downstream sequence were linked together. The resulting fragment was ligated into the pMD-18T vector to generate a donor plasmid for | this study |

| | | | | |
|---|---|---|---|---|
| | | | insertion of the *EDEM2* cassette into the Sg19 locus of FIM1ΔU, KMU1&2-S and KMU1&2-S-E110A. | |
| LHZ2036 | *AmpR*, Sg19 upstream-$P_{ScTEF1}$-$SP_{KmPDI1}$-*HsEDEM2*-*E117Q-HDEL*-$T_{ScTEF1}$-Sg19 downstream | pMD-18T | All elements were identical to those in LHZ2035, except that the *HsEDEM2* CDS carried the E117Q substitution. The resulting fragment was ligated into the pMD-18T vector to generate a donor plasmid for insertion of the *EDEM2-E117Q* cassette into the Sg19 locus of FIM1ΔU. | this study |
| LHZ2037 | *AmpR*, Sg7 upstream-$P_{ScADH2}$ -$SP_{KmCNE1}$-*HsCANX*-$T_{ScADH2}$-Sg7 downstream | pMD-18T | 500 bp Sg7 upstream sequence, *ScADH2*[1] promoter[2] (Gene ID 855349), *KmCNE1* signal sequence (Gene ID 34716631, 1-54 bp), CDS of *HsCANX* (Gene ID: 821, 63–1776 bp), 1000 bp *ScADH2* terminator[3], 500 bp Sg7 downstream sequence were linked together. The resulting fragment was ligated into the pMD-18T vector to generate a donor plasmid for insertion of the *CANX* cassette into the Sg7 locus of FIM1ΔU. | this study |
| LHZ2038 | *AmpR*, Sg7 upstream-$P_{ScTEF2}$-$SP_{KmPDI1}$-*HsCALR*-*HDEL*-$T_{ScTEF2}$-Sg7 downstream | pMD-18T | 500 bp Sg7 upstream sequence, *ScTEF2*[1] promoter[2] (Gene ID 852415), *KmPDI1* signal sequence, CDS of *HsCALR* (Gene ID: 811, 54–1251 bp), 1000 bp *ScTEF2* terminator[3], 500 bp Sg7 downstream sequence were linked together. The resulting fragment was ligated into the pMD-18T vector to generate | this study |

| | | | | |
|---|---|---|---|---|
| | | | a donor plasmid for insertion of the *CALR* cassette into the Sg7 locus of FIM1ΔU. | |
| LHZ2039 | *AmpR*, Sg7 upstream-P$_{ScPGK1}$-SP$_{KmPDI1}$-*HsPDIA3-HDEL*-T$_{ScPGK1}$-Sg7 downstream | pMD-18T | 500 bp Sg7 upstream sequence, *ScPGK1*[1] promoter[2] (Gene ID 850370), *KmPDI1* signal sequence, CDS of *HsPDIA3* (Gene ID: 2923, 75–1515 bp), 1000 bp *ScPGK1* terminator[3], 500 bp Sg7 downstream sequence were linked together. The resulting fragment was ligated into the pMD-18T vector to generate a donor plasmid for insertion of the *PDIA3* cassette into the Sg7 locus of FIM1ΔU. | this study |
| LHZ2040 | *AmpR*, Sg7 upstream-P$_{ScTDH3}$-SP$_{KmPDI1}$-*HsPPIB-HDEL*-T$_{ScTDH3}$-P$_{PpAFT1}$-SP$_{KmPDI1}$-*HsERP29-HDEL*-T$_{PpAFT1}$-Sg7 downstream | pMD-18T | 500 bp Sg7 upstream sequence, *ScTDH3*[1] promoter[2] (Gene ID 853106), *KmPDI1* signal sequence, CDS of *HsPPIB* (Gene ID: 5479, 102–648 bp), 1000 bp *ScTDH3* terminator[3], were linked together. *PpAFT1*[1] promoter[2] (Gene ID 8196669), *KmPDI1* signal sequence, CDS of *HsERP29* (Gene ID: 10961, 99–783 bp), 1000 bp *PpAFT1* terminator[3], 500 bp Sg7 downstream sequence were linked together. The *HsPPIB* and *HsERP29* expression cassettes were subsequently fused to obtain the final fragment. The resulting fragment was ligated into the pMD-18T vector to generate a donor plasmid for insertion of the *PPIB* and *ERP29* cassettes into the Sg7 locus of FIM1ΔU. | this study |

| | | | | |
|---|---|---|---|---|
| LHZ2041 | *AmpR*, Sg7 upstream-$P_{ScPGK1}$-$SP_{KmPDI1}$-*HsPDIA3-HDEL*-$T_{ScPGK1}$- N-terminal $P_{ScADH2}$ | pMD-18T | 500 bp Sg7 upstream sequence, *ScPGK1*[1] promoter[2] (Gene ID 850370), *KmPDI1* signal sequence, CDS of *HsPDIA3* (Gene ID: 2923, 75–1515 bp), 1000 bp *ScPGK1* terminator[3], a 500-bp homologous sequence corresponding to the N-terminal sequence of $P_{ScADH2}$ promoter in the *CANX* cassette were linked together. The resulting fragment was ligated into the pMD-18T vector to generate a donor plasmid for insertion of the *PDIA3* cassette upstream of the *CANX* cassette in $KM_{CNX}$. | this study |
| LHZ2042 | *AmpR*, Sg7 upstream-$P_{ScPGK1}$-$SP_{KmPDI1}$-*HsPDIA3-HDEL*-$T_{ScPGK1}$- N-terminal $P_{ScTEF2}$ | pMD-18T | 500 bp Sg7 upstream sequence, *ScPGK1*[1] promoter[2] (Gene ID 850370), *KmPDI1* signal sequence, CDS of *HsPDIA3* (Gene ID: 2923, 75–1515 bp), 1000 bp *ScPGK1* terminator[3], a 500-bp homologous sequence corresponding to the N-terminal sequence of the $P_{ScTEF2}$ in the *CALR* cassette were linked together. The resulting fragment was ligated into the pMD-18T vector to generate a donor plasmid for insertion of the *PDIA3* cassette upstream of the *CALR* cassette in $KM_{CRT}$. | this study |
| LHZ2043 | *AmpR*, Sg7 upstream-$P_{ScTDH3}$-$SP_{KmPDI1}$-*HsPPIB*-*HDEL*-$T_{ScTDH3}$-$P_{PpAFT1}$-$SP_{KmPDI1}$- | pMD-18T | 500 bp Sg7 upstream sequence, *ScTDH3*[1] promoter[2] (Gene ID 853106), *KmPDI1* signal sequence, CDS of *HsPPIB* (Gene ID: 5479, 102–648 bp), 1000 bp *ScTDH3* terminator[3] were linked together. *PpAFT1*[1] promoter[2] | this study |

| | | | | |
|---|---|---|---|---|
| | *HsERP29-HDEL*-$T_{PpAFT1}$- N-terminal $P_{ScPGK1}$ | | (Gene ID 8196669), *KmPDI1* signal sequence, CDS of *HsERP29* (Gene ID: 10961, 99–783 bp), 1000 bp *PpAFT1* terminator[3], a 500 bp homologous sequence corresponding to the N-terminal sequence of $P_{ScPGK1}$ in the *PDIA3* cassette were linked together. The *HsPPIB* and *HsERP29* expression cassettes were subsequently fused to obtain the final fragment. The resulting fragment was ligated into the pMD-18T vector to generate a donor plasmid for insertion of the *PPIB* and *ERP29* cassettes upstream of the *PDIA3* cassette in $KM_{CNX-p57}$. | |
| LHZ2044 | *AmpR*, Sg7 upstream-$P_{ScTDH3}$-$SP_{KmPDI1}$-*HsPPIB*-*HDEL*-$T_{ScTDH3}$-$P_{PpAFT1}$-$SP_{KmPDI1}$-*HsERP29-HDEL*-$T_{PpAFT1}$- N-terminal $P_{ScPGK1}$ | pMD-18T | 500 bp Sg7 upstream sequence, *ScTDH3*[1] promoter[2] (Gene ID 853106), *KmPDI1* signal sequence, CDS of *HsPPIB* (Gene ID: 5479, 102–648 bp), 1000 bp *ScTDH3* terminator[3] were linked together. *PpAFT1*[1] promoter[2] (Gene ID 8196669), *KmPDI1* signal sequence, CDS of *HsERP29* (Gene ID: 10961, 99–783 bp), 1000 bp *PpAFT1* terminator[3], a 500 bp homologous sequence corresponding to the N-terminal sequence of $P_{ScPGK1}$ in the *PDIA3* cassette were linked together. The *HsPPIB* and *HsERP29* expression cassettes were subsequently fused to obtain the final fragment. The resulting fragment was ligated into the pMD- | this study |

| | | | | |
|---|---|---|---|---|
| | | | 18T vector to generate a donor plasmid for insertion of the *PPIB* and *ERP29* cassettes upstream of the PDIA3 cassette in $KM_{CRT\text{-}p57}$. | |
| LHZ2045 | *AmpR*, *KmGTB1* upstream-*KmGTB1* downstream | pMD-18T | 500 bp upstream and downstream sequences flanking the *KmGTB1* (Gene ID: 34714607) were linked together. The resulting fragment was ligated into the pMD-18T vector to generate a donor plasmid for deletion of *GTB1* in FIM1ΔU or $KM_{U1\&2\text{-}S}$. | this study |
| LHZ2046 | *AmpR*, *KmGTB1* upstream-*KmGTB1-E110A* *KmGTB1* downstream | pMD-18T | 500 bp *KmGTB1* (Gene ID: 34714607) upstream sequence, the *KmGTB1* coding sequence carrying the E110A substitution, and 500 bp *KmGTB1* downstream sequence were linked together. The resulting fragment was ligated into the pMD-18T vector to generate a donor plasmid for integration of the *GTB1-E110A* into the *KmGTB1* locus in $KM_{\mathit{gtb1}\Delta}$ or $KM_{U1\&2\text{-}S\text{-}\mathit{gtb1}\Delta}$. | this study |
| LHZ2047 | *AmpR*, *KmGTB1* upstream-*KmGTB1-R629A* *KmGTB1* downstream | pMD-18T | 500 bp *KmGTB1* (Gene ID: 34714607) upstream sequence, the *KmGTB1* coding sequence carrying the R629A substitution, and 500 bp *KmGTB1* downstream sequence were linked together. The resulting fragment was ligated into the pMD-18T vector to generate a donor plasmid for integration of the *GTB1-R629A* into the *KmGTB1* locus in $KM_{\mathit{gtb1}\Delta}$. | this study |

| | | | | |
|---|---|---|---|---|
| LHZ2048 | *AmpR*, *KmGTB1* upstream-*KmGTB1-Y654F KmGTB1* downstream | pMD-18T | 500 bp *KmGTB1* (Gene ID: 34714607) upstream sequence, the *KmGTB1* coding sequence carrying the Y654F substitution, and 500 bp *KmGTB1* downstream sequence were linked together. The resulting fragment was ligated into the pMD-18T vector to generate a donor plasmid for integration of the *GTB1-Y654F* into the *KmGTB1* locus in $KM_{gtb1\Delta}$. | this study |
| LHZ2049 | *AmpR*, *KmCNE1* upstream-*KmCNE1* downstream | pMD-18T | 500 bp upstream and downstream sequences flanking the *KmCNE1* (Gene ID: 34716631) were linked together. The resulting fragment was ligated into the pMD-18T vector to generate a donor plasmid for deletion of the *CNE1* in FIM1ΔU or $KM_{UGGT1}$. | this study |
| LHZ2050 | *AmpR*, *KmROT2* upstream-*KmROT2* downstream | pMD-18T | 500 bp upstream and downstream sequences flanking the *KmROT2* (Gene ID: 34714747) were linked together. The resulting fragment was ligated into the pMD-18T vector to generate a donor plasmid for deletion of the *ROT2* in FIM1ΔU or $KM_{gtb1\Delta}$. | this study |
| LHZ2051 | *AmpR*, *KmROT2* upstream-$SP_{KmPDI1}$-*HsGANAB-HDEL-KmROT2* downstream | pMD-18T | 500 bp *KmROT2* upstream sequence, *KmPDI1* signal sequence, CDS of *HsGANAB* (Gene ID: 23193, 87–2832 bp), 500 bp *KmROT2* downstream sequence were linked together. The resulting fragment was ligated into the pMD-18T vector to generate a donor plasmid for insertion of the | this study |

| | | | | |
|---|---|---|---|---|
| | | | *GANAB* fragment into the *KmROT2* locus in KM$_{rot2Δ}$. | |
| LHZ2052 | *AmpR*, *KmGTB1* upstream-SP$_{KmPDI1}$-*HsPRKCSH*-*HDEL*- *KmGTB1* downstream | pMD-18T | 500 bp *KmGTB1* upstream sequence, *KmPDI1* signal sequence, CDS of *HsPRKCSH* (Gene ID: 5589, 45–1584 bp), 500 bp *KmGTB1* downstream sequence were linked together. The resulting fragment was ligated into the pMD-18T vector to generate a donor plasmid for insertion of the *PRKCSH* fragment into the *KmGTB1* locus in KM$_{hGII\alpha}$. | this study |
| LHZ2053 | *AmpR*, *KmMNS1* upstream-*KmMNS1* downstream | pMD-18T | 500 bp upstream and downstream sequences flanking the *KmMNS1* (Gene ID: 34716495) were linked together. The resulting fragment was ligated into the pMD-18T vector to generate a donor plasmid for deletion of the *MNS1* in FIM1ΔU. | this study |
| LHZ531 | *KmARS1*/*CEN5*, *KmURA3*, SpCas9, gRNA | / | Backbone vector for SpCas9 plasmids | [1] |
| LHZ2054 | *KmARS1*/*CEN5*, *KmURA3*, SpCas9, gRNA | LHZ531 | gRNA-302F/gRNA-302R primer pair inserted into SapI sites of LHZ531, generating a CRISPR plasmid targeting Sg7 locus. Used for insertion of the *SEP15-U96C*-cassette in FIM1ΔU, KM$_{UGGT1}$, KM$_{UGGT2}$ or KM$_{U1\&2}$. Also for insertion of the *CANX* cassette, *CALR* cassette, *PDIA3* cassette, *PPIB* and *ERP29* cassette in FIM1ΔU. | this study |

| | | | | |
|---|---|---|---|---|
| LHZ2055 | *KmARS1*/*CEN5*, *KmURA3*, SpCas9, gRNA | LHZ531 | gRNA-302-CANXF/gRNA-302-CANXR primer pair inserted into SapI sites of LHZ531, generating a CRISPR plasmid targeting $P_{ScADH2}$ of *CANX* cassette. Used for insertion of the *PDIA3* cassette in $KM_{CNX}$. | this study |
| LHZ2056 | *KmARS1*/*CEN5*, *KmURA3*, SpCas9, gRNA | LHZ531 | gRNA-302-CALRF/gRNA-302-CALRR primer pair inserted into SapI sites of LHZ531, generating a CRISPR plasmid targeting $P_{ScTEF2}$ of *CALR* cassette. Used for insertion of the *PDIA3* cassette in $KM_{CRT}$. | this study |
| LHZ2057 | *KmARS1*/*CEN5*, *KmURA3*, SpCas9, gRNA | LHZ531 | gRNA-302-2CF/gRNA-302-2CR primer pair inserted into SapI sites of LHZ531, generating a CRISPR plasmid targeting $P_{ScPGK1}$ of *PDIA3* cassette. Used for insertion of the *PPIB* and *ERP29* cassette in $KM_{CNX-p57}$ or $KM_{CRT-p57}$. | this study |
| LHZ2058 | *KmARS1*/*CEN5*, *KmURA3*, SpCas9, gRNA | LHZ531 | gRNA-702F/gRNA-702R primer pair inserted into SapI sites of LHZ531, generating a CRISPR plasmid targeting Sg19 locus. Used for insertion of the *EDEM2* cassette in FIM1ΔU, $KM_{U1\&2-S}$, and $KM_{U1\&2-S-E110A}$; and for insertion of *EDEM2-E117Q* in FIM1ΔU. | this study |
| LHZ2059 | *KmARS1*/*CEN5*, *KmURA3*, SpCas9, gRNA | LHZ531 | gRNA-GTB1-KO-F/gRNA-GTB1-KO-R primer pair inserted into SapI sites of LHZ531, generating a CRISPR plasmid targeting 3′ end of *GTB1* locus. Used for deleting *GTB1* in FIM1ΔU or $KM_{U1\&2-S}$. | this study |

| | | | | |
|---|---|---|---|---|
| LHZ2060 | *KmARS1*/*CEN5*, *KmURA3*, SpCas9, gRNA | LHZ531 | gRNA-ROT2-KO-F/gRNA-ROT2-KO-R primer pair inserted into SapI sites of LHZ531, generating a CRISPR plasmid targeting 3′ end of *ROT2* locus. Used for deleting *ROT2* in FIM1ΔU or $KM_{gtb1Δ}$. | this study |
| LHZ2061 | *KmARS1*/*CEN5*, *KmURA3*, SpCas9, gRNA | LHZ531 | gRNA-MNS1-KO-F/gRNA-MNS1-KO-R primer pair inserted into SapI sites of LHZ531, generating a CRISPR plasmid targeting 3′ end of *MNS1* locus. Used for deleting *MNS1* in FIM1ΔU. | this study |
| LHZ2062 | *KmARS1*/*CEN5*, *KmURA3*, SpCas9, gRNA | LHZ531 | gRNA-CNE1-KO-F/gRNA-CNE1-KO-R primer pair inserted into SapI sites of LHZ531, generating a CRISPR plasmid targeting 3′ end of *CNE1* locus. Used for deleting *CNE1* in FIM1ΔU or $KM_{UGGT1}$. | this study |
| LHZ2063 | *KmARS1*/*CEN5*, *KmURA3*, SpCas9, gRNA | LHZ531 | gRNA-GTB1-KI-F/gRNA-GTB1-KI-R primer pair inserted into SapI sites of LHZ531, generating a CRISPR plasmid targeting a site in the 3′ flanking region of *GTB1*, immediately downstream of the stop codon. Used for insertion of the *PRKCSH* cassette, in $KM_{hGII\alpha}$, and for insertion of GTB1-*E110A* cassette, GTB1-*R629A* cassette and GTB1-*Y654F* cassette in $KM_{gtb1Δ}$ or $KM_{U1\&2\text{-}S\text{-}gtb1Δ}$. | this study |
| LHZ2064 | *KmARS1*/*CEN5*, *KmURA3*, SpCas9, gRNA | LHZ531 | gRNA-ROT2-KI-F/gRNA-ROT2-KI-R primer pair inserted into SapI sites of LHZ531, generating a CRISPR plasmid targeting a site in the 3′ flanking region of *ROT2*, | this study |

| | | | | |
|---|---|---|---|---|
| | | | immediately downstream of the stop codon. Used for insertion of the *GANAB* cassette in $KM_{rot2\Delta}$. | |
| LHZ2065 | *KmARS1*/*CEN5*, *KmURA3*, Sta2Cas9, gRNA | LHZ531 | Backbone vector for Sta2-Cas9 plasmids | this study |
| LHZ2066 | *KmARS1*/*CEN5*, *KmURA3*, Sta2Cas9, gRNA | LHZ2065 | gRNA-UGGT1F gRNA-UGGT1R primer pair inserted into AarI sites of LHZ2065, generating a CRISPR plasmid targeting Sg1 locus. Used for insertion of the *UGGT1* cassette or *UGGT1-D1358A* in FIM1ΔU. | this study |
| LHZ2067 | *KmARS1*/*CEN5*, *KmURA3*, Sta2Cas9, gRNA | LHZ2065 | gRNA-UGGT2F gRNA-UGGT2R primer pair inserted into AarI sites of LHZ2065, generating a CRISPR plasmid targeting Sg4 locus. Used for insertion of the *UGGT2* cassette in FIM1ΔU or $KM_{UGGT1}$ or *UGGT2-D1333A* in FIM1ΔU. | this study |
| LHZ2068 | pKD1, $P_{KmINU1}$-$SP_{KmINU1}$-Fc-$His_6$-$T_{KmINU1}$, *URA3* | LHZ399 | Secretory expression of human Fc (Gene ID 3500) fragment, 6-His tag was added to the C terminal. The Fc coding sequence was codon-optimized based on the human IgG1 Fc region derived from IGHG1. The sequence can be found in Table S3. | this study |
| LHZ2069 | pKD1, $P_{KmINU1}$-$SP_{KmINU1}$-GGH-$His_6$-$T_{KmINU1}$, *URA3* | LHZ399 | Secretory expression of human GGH (Gene ID 8836. 75-954 bp), 6-His tag was added to the C terminal. | this study |

| | | | | |
|---|---|---|---|---|
| LHZ443 | pKD1, $P_{KmINU1}$-$SP_{KmINU1}$-Xyn-$T_{KmINU1}$, *URA3* | / | Secretory expression of Xyn-CDBFV | Ref. (2018 BB) |
| LHZ2070 | pKD1, $P_{KmINU1}$-$SP_{KmINU1}$-dulaglutide-$T_{KmINU1}$, *URA3* | LHZ399 | Secretory expression of dulaglutide (DrugBank ID DB09045). | this study |
| LHZ2071 | pKD1, $P_{KmINU1}$-$SP_{KmINU1}$-abatacept-$T_{KmINU1}$, *URA3* | LHZ399 | Secretory expression of abatacept (DrugBank ID DB01281). | this study |
| LHZ2072 | pKD1, $P_{KmINU1}$-$SP_{KmINU1}$-etanercept-$T_{KmINU1}$, *URA3* | LHZ399 | Secretory expression of etanercept (DrugBank ID DB00005). | this study |
| LHZ399 | pKD1, $P_{KmINU1}$-$SP_{KmINU1}$-filler fragment-$T_{KmINU1}$, *URA3* | / | Backbone plasmid containing the INU expression cassette | [2] |
| pUKDN132 | pKD1, $P_{KmINU1}$-$SP_{KmINU1}$-$T_{KmINU1}$, *URA3* | / | empty vector | [2] |

[1] Species abbreviation: *Sc*, *Saccharomyces cerevisiae*; Pp: *Pichia pastoris*; *Km*, *Kluyveromyces marxianus*; *Hs*, *Homo sapiens*.

[2] The promoter is defined as the 1000 bp sequence upstream of the start codon.

[3]. The terminator is defined as the sequence downstream of the stop codon.

[1] ZHOU H, TIAN T, LIU J, et al. Efficient and markerless gene integration with SlugCas9-HF in Kluyveromyces marxianus [J]. Commun Biol, 2024, 7(1): 797.

[2] ZHOU J, ZHU P, HU X, et al. Improved secretory expression of lignocellulolytic enzymes in Kluyveromyces marxianus by promoter and signal sequence engineering [J]. Biotechnol Biofuels, 2018, 11: 235.
