## Supplementary Table 2. Strains used in this study for "Humanization of N-glycan-dependent protein quality control system in *Kluyveromyces marxianus* promotes glycoprotein secretion"

Table S2. Strains used in this study.

| Name | Genotype | Construction details：CRISPR plasmid, donor plasmid, host strain | Source or reference |
|---|---|---|---|
| FIM1ΔU | *ura3*Δ | / | [1] |
| KM$_{UGGT1}$ | *ura3*Δ<br>Sg1::P$_{ScADH1}$-SP$_{KmPDI1}$-*UGGT1*-T$_{ScADH1}$ | LHZ2066, LHZ2030, FIM1ΔU | this study |
| KM$_{UGGT1-D1358A}$ | *ura3*Δ<br>Sg1::P$_{ScADH1}$-SP$_{KmPDI1}$-*UGGT1-D1358A*-T$_{ScADH1}$ | LHZ2066, LHZ2031, FIM1ΔU | this study |
| KM$_{UGGT2}$ | *ura3*Δ<br>Sg4::P$_{ScGPD1}$-SP$_{KmPDI1}$-*UGGT2*-T$_{ScGPD1}$ | LHZ2067, LHZ2032, FIM1ΔU | this study |
| KM$_{U2-D1333A}$ | *ura3*Δ<br>Sg4::P$_{ScGPD1}$-SP$_{KmPDI1}$-*UGGT2-D1333A* -T$_{ScGPD1}$ | LHZ2067, LHZ2033, FIM1ΔU | this study |
| KM$_{U1\&2}$ | *ura3*Δ<br>Sg1::P$_{ScADH1}$-SP$_{KmPDI1}$-*UGGT1*-T$_{ScADH1}$<br>Sg4::P$_{ScGPD1}$-SP$_{KmPDI1}$-*UGGT2*-T$_{ScGPD1}$ | LHZ2067, LHZ2032, KM$_{UGGT1}$ | this study |
| KM$_{SEP15}$ | *ura3*Δ<br>Sg7::P$_{ScCYS3}$-SP$_{KmPDI1}$-*SEP15*-U96C-T$_{ScCYS3}$ | LHZ2054, LHZ2034, FIM1ΔU | this study |
| KM$_{U1-S}$ | *ura3*Δ<br>Sg1::P$_{ScADH1}$-SP$_{KmPDI1}$-*UGGT1*-T$_{ScADH1}$<br>Sg7::P$_{ScCYS3}$-SP$_{KmPDI1}$-*SEP15*-U96C-T$_{ScCYS3}$ | LHZ2054, LHZ2034, KM$_{UGGT1}$ | this study |
| KM$_{U2-S}$ | *ura3*Δ<br>Sg4::P$_{ScGPD1}$-SP$_{KmPDI1}$-*UGGT2*-T$_{ScGPD1}$<br>Sg7::P$_{ScCYS3}$-SP$_{KmPDI1}$-*SEP15*-U96C-T$_{ScCYS3}$ | LHZ2054, LHZ2034, KM$_{UGGT2}$ | this study |
| KM$_{U1\&2-S}$ | *ura3*Δ | LHZ2054, LHZ2034, KM$_{U1\&2}$ | this study |

| | | | |
|---|---|---|---|
| | Sg1::P$_{ScADH1}$-SP$_{KmPDI1}$-*UGGT1*-T$_{ScADH1}$<br>Sg4::P$_{ScGPD1}$-SP$_{KmPDI1}$-*UGGT2*-T$_{ScGPD1}$<br>Sg7::P$_{ScCYS3}$-SP$_{KmPDI1}$-*SEP15*-U96C-T$_{ScCYS3}$ | | |
| KM$_{cne1\Delta}$ | *ura3*Δ *cne1*Δ | LHZ2062, LHZ2049, FIM1ΔU | this study |
| KM$_{U1\text{-}cne1\Delta}$ | *ura3*Δ<br>*cne1*Δ<br>Sg1::P$_{ScADH1}$-SP$_{KmPDI1}$-*UGGT1*-T$_{ScADH1}$ | LHZ2062, LHZ2049, KM$_{UGGT1}$ | this study |
| KM$_{CNX}$ | *ura3*Δ<br>Sg7::P$_{ScADH2}$-SP$_{KmCNE1}$-CANX-T$_{ScADH2}$ | LHZ2054, LHZ2037, FIM1ΔU | this study |
| KM$_{CRT}$ | *ura3*Δ<br>Sg7::P$_{ScTEF2}$-SP$_{KmPDI1}$-*CALR*-T$_{ScTEF2}$ | LHZ2054, LHZ2038, FIM1ΔU | this study |
| KM$_{p57}$ | *ura3*Δ<br>Sg7::P$_{ScPGK1}$-SP$_{KmPDI1}$-*PDIA3*-T$_{ScPGK1}$ | LHZ2054, LHZ2039, FIM1ΔU | this study |
| KM$_{cypb\text{-}p29}$ | *ura3*Δ<br>Sg7:: P$_{ScTDH3}$-SP$_{KmPDI1}$-*PPIB*-T$_{ScTDH3}$-<br>P$_{PpAFT1}$-SP$_{KmPDI1}$-*ERP29*-T$_{PpAFT1}$- | LHZ2054, LHZ2040, FIM1ΔU | this study |
| KM$_{CNX\text{-}p57}$ | *ura3*Δ<br>Sg7::P$_{ScPGK1}$-SP$_{KmPDI1}$-*PDIA3*-T$_{ScPGK1}$-<br>P$_{ScADH2}$-SP$_{KmCNE1}$-CANX-T$_{ScADH2}$ | LHZ2055, LHZ2041, KM$_{CNX}$ | this study |
| KM$_{U1\text{-}CNX\text{-}p57}$ | *ura3*Δ<br>Sg1::P$_{ScADH1}$-SP$_{KmPDI1}$-*UGGT1*-T$_{ScADH1}$<br>Sg7::P$_{ScPGK1}$-SP$_{KmPDI1}$-*PDIA3*-T$_{ScPGK1}$-<br>P$_{ScADH2}$-SP$_{KmCNE1}$-CANX-T$_{ScADH2}$ | LHZ2066, LHZ2030, KM$_{CNX\text{-}p57}$ | this study |
| KM$_{CRT\text{-}p57}$ | *ura3*Δ<br>Sg7::P$_{ScPGK1}$-SP$_{KmPDI1}$-*PDIA3*-T$_{ScPGK1}$-<br>P$_{ScTEF2}$-SP$_{KmPDI1}$-*CALR*-T$_{ScTEF2}$ | LHZ2056, LHZ2042, KM$_{CRT}$ | this study |

| | | | |
|---|---|---|---|
| KM$_{U1-CRT-p57}$ | *ura3*Δ<br>Sg1::P$_{ScADH1}$-SP$_{KmPDI1}$-*UGGT1*-T$_{ScADH1}$<br>Sg7::P$_{ScPGK1}$-SP$_{KmPDI1}$-*PDIA3*-T$_{ScPGK1}$-<br>P$_{ScTEF2}$-SP$_{KmPDI1}$-*CALR*-T$_{ScTEF2}$ | LHZ2066, LHZ2030, KM$_{CRT-p57}$ | this study |
| KM$_{CNX-3C}$ | *ura3*Δ<br>Sg7:: P$_{ScTDH3}$-SP$_{KmPDI1}$-*PPIB*-T$_{ScTDH3}$-<br>P$_{PpAFT1}$-SP$_{KmPDI1}$-*ERP29*-T$_{PpAFT1}$-<br>P$_{ScPGK1}$-SP$_{KmPDI1}$-*PDIA3*-T$_{ScPGK1}$-<br>P$_{ScADH2}$-SP$_{KmCNE1}$-*CANX*-T$_{ScADH2}$ | LHZ2057, LHZ2043, KM$_{CNX-p57}$ | this study |
| KM$_{CRT-3C}$ | *ura3*Δ<br>Sg7:: P$_{ScTDH3}$-SP$_{KmPDI1}$-*PPIB*-T$_{ScTDH3}$-<br>P$_{PpAFT1}$-SP$_{KmPDI1}$-*ERP29*-T$_{PpAFT1}$-<br>P$_{ScPGK1}$-SP$_{KmPDI1}$-*PDIA3*-T$_{ScPGK1}$-<br>P$_{ScTEF2}$-SP$_{KmPDI1}$-*CALR*-T$_{ScTEF2}$ | LHZ2057, LHZ2044, KM$_{CRT-p57}$ | this study |
| KM$_{U1-CNX-3C}$ | *ura3*Δ<br>Sg1::P$_{ScADH1}$-SP$_{KmPDI1}$-*UGGT1*-T$_{ScADH1}$<br>Sg7:: P$_{ScTDH3}$-SP$_{KmPDI1}$-*PPIB*-T$_{ScTDH3}$-<br>P$_{PpAFT1}$-SP$_{KmPDI1}$-*ERP29*-T$_{PpAFT1}$-<br>P$_{ScPGK1}$-SP$_{KmPDI1}$-*PDIA3*-T$_{ScPGK1}$-<br>P$_{ScADH2}$-SP$_{KmCNE1}$-*CANX*-T$_{ScADH2}$ | LHZ2066, LHZ2030, KM$_{CNX-3C}$ | this study |
| KM$_{U1-CRT-3C}$ | *ura3*Δ<br>Sg1::P$_{ScADH1}$-SP$_{KmPDI1}$-*UGGT1*-T$_{ScADH1}$<br>Sg7:: P$_{ScTDH3}$-SP$_{KmPDI1}$-*PPIB*-T$_{ScTDH3}$-<br>P$_{PpAFT1}$-SP$_{KmPDI1}$-*ERP29*-T$_{PpAFT1}$-<br>P$_{ScPGK1}$-SP$_{KmPDI1}$-*PDIA3*-T$_{ScPGK1}$-<br>P$_{ScTEF2}$-SP$_{KmPDI1}$-*CALR*-T$_{ScTEF2}$ | LHZ2066, LHZ2030, KM$_{CRT-3C}$ | this study |
| KM$_{rot2Δ}$ | *ura3*Δ *rot2*Δ | LHZ2060, LHZ2050, FIM1ΔU | this study |

| | | | |
|---|---|---|---|
| $KM_{gtb1\Delta}$ | *ura3*Δ *gtb1*Δ | LHZ2059, LHZ2045, FIM1ΔU | this study |
| $KM_{GII\Delta}$ | *ura3*Δ *rot2*Δ *gtb1*Δ | LHZ2060, LHZ2050, $KM_{gtb1\Delta}$ | this study |
| $KM_{hGII\alpha}$ | *ura3*Δ<br>*rot2*Δ::*GANAB* | LHZ2064, LHZ2051, $KM_{rot2\Delta}$ | this study |
| $KM_{hGII}$ | *ura3*Δ<br>*rot2*Δ::*GANAB*<br>*gtb1*Δ::*PRKCSH* | LHZ2063, LHZ2052, $KM_{hGII\alpha}$ | this study |
| $KM_{U1\text{-}hGII}$ | *ura3*Δ<br>*rot2*Δ::*GANAB*<br>*gtb1*Δ::*PRKCSH*<br>Sg1::$P_{ScADH1}$-$SP_{KmPDI1}$-*UGGT1*-$T_{ScADH1}$ | LHZ2066, LHZ2030, $KM_{hGII}$ | this study |
| $KM_{\beta\text{-}E110A}$ | *ura3*Δ<br>*gtb1*Δ::*GTB1-E110A* | LHZ2063, LHZ2046, $KM_{gtb1\Delta}$ | this study |
| $KM_{\beta\text{-}R629A}$ | *ura3*Δ<br>*gtb1*Δ::*GTB1-R629A* | LHZ2063, LHZ2047, $KM_{gtb1\Delta}$ | this study |
| $KM_{\beta\text{-}Y654F}$ | *ura3*Δ<br>*gtb1*Δ::*GTB1-Y654F* | LHZ2063, LHZ2048, $KM_{gtb1\Delta}$ | this study |
| $KM_{EDEM2}$ | *ura3*Δ<br>Sg19::$P_{ScTEF1}$-$SP_{KmPDI1}$-*EDEM2*-$T_{ScTEF1}$ | LHZ2058, LHZ2035, FIM1ΔU | this study |
| $KM_{E\text{-}E117Q}$ | *ura3*Δ<br>Sg19::$P_{ScTEF1}$-$SP_{KmPDI1}$-*EDEM2-E117Q*-$T_{ScTEF1}$ | LHZ2058, LHZ2036, FIM1ΔU | this study |
| $KM_{mns1\Delta}$ | *ura3*Δ *mns1*Δ | LHZ2061, LHZ2053, FIM1ΔU | this study |
| $KM_{U1\&2\text{-}S\text{-}gtb1\Delta}$ | *ura3*Δ<br>Sg1::$P_{ScADH1}$-$SP_{KmPDI1}$-*UGGT1*-$T_{ScADH1}$ | LHZ2059, LHZ2045, $KM_{U1\&2\text{-}S}$ | this study |

| | | | |
|---|---|---|---|
| | Sg4::P$_{ScGPD1}$-SP$_{KmPDI1}$-*UGGT2*-T$_{ScGPD1}$<br>Sg7::P$_{ScCYS3}$-SP$_{KmPDI1}$-*SEP15*-U96C-T$_{ScCYS3}$<br>*gtb1*Δ | | |
| KM$_{U1\&2-S-E110A}$ | *ura3*Δ<br>Sg1::P$_{ScADH1}$-SP$_{KmPDI1}$-*UGGT1*-T$_{ScADH1}$<br>Sg4::P$_{ScGPD1}$-SP$_{KmPDI1}$-*UGGT2*-T$_{ScGPD1}$<br>Sg7::P$_{ScCYS3}$-SP$_{KmPDI1}$-*SEP15*-U96C-T$_{ScCYS3}$<br>*gtb1*Δ::*GTB1-E110A* | LHZ2063, LHZ2046, KM$_{U1\&2-S-gtb1\Delta}$ | this study |
| KM$_{U1\&2-S-EDEM2}$ | *ura3*Δ<br>Sg1::P$_{ScADH1}$-SP$_{KmPDI1}$-*UGGT1*-T$_{ScADH1}$<br>Sg4::P$_{ScGPD1}$-SP$_{KmPDI1}$-*UGGT2*-T$_{ScGPD1}$<br>Sg7::P$_{ScCYS3}$-SP$_{KmPDI1}$-*SEP15*-U96C-T$_{ScCYS3}$<br>Sg19::P$_{ScTEF1}$-SP$_{KmPDI1}$-*EDEM2*-T$_{ScTEF1}$ | LHZ2058, LHZ2035, KM$_{U1\&2-S}$ | this study |
| KM$_{ultra}$ | *ura3*Δ<br>Sg1::P$_{ScADH1}$-SP$_{KmPDI1}$-*UGGT1*-T$_{ScADH1}$<br>Sg4::P$_{ScGPD1}$-SP$_{KmPDI1}$-*UGGT2*-T$_{ScGPD1}$<br>Sg7::P$_{ScCYS3}$-SP$_{KmPDI1}$-*SEP15*-U96C-T$_{ScCYS3}$<br>Sg19::P$_{ScTEF1}$-SP$_{KmPDI1}$-*EDEM2*-T$_{ScTEF1}$<br>*gtb1*Δ::*GTB1-E110A* | LHZ2058, LHZ2035, KM$_{U1\&2-S-E110A}$ | this study |


1. ZHOU J, ZHU P, HU X, et al. Improved secretory expression of lignocellulolytic enzymes in Kluyveromyces marxianus by promoter and signal sequence engineering [J]. Biotechnol Biofuels, 2018, 11: 235.
