## Supplementary Table 3. Primers and synthetic DNA sequences used in this study for "Humanization of N-glycan-dependent protein quality control system in *Kluyveromyces marxianus* promotes glycoprotein secretion"

Table S3. Primers and synthetic DNA sequences used in this study.

| Name | Sequence | Details |
|---|---|---|
| gRNA-UGGT1F | ATTGGGGGAATTATAAGCGTA | Sta2-Cas9 gRNA, Target Sg1; used for integration of *UGGT1* or *UGGT1-D1358A* cassette |
| gRNA-UGGT1R | TACGCTTATAATTCCCCCAAT | Sta2-Cas9 gRNA, Target Sg1; used for integration of *UGGT1* or *UGGT1-D1358A* cassette |
| gRNA-UGGT2F | TATATTTCCTTCCTTCCACTA | Sta2-Cas9 gRNA, Target Sg4; used for integration of *UGGT2* or *UGGT2-D1333A* cassette |
| gRNA-UGGT2R | TAGTGGAAGGAAGGAAATATA | Sta2-Cas9 gRNA, Target Sg4; used for integration of *UGGT2* or *UGGT2-D1333A* cassette |
| gRNA-302F | ACACAGCATTGGAAATACCA | Target Sg7; used for *SEP15*, *CANX*, *CALR*, *PDIA3*, *PPIB-ERP29* insertion at Sg7 |
| gRNA-302R | TGGTATTTCCAATGCTGTGT | Target Sg7; used for *SEP15*, *CANX*, *CALR*, *PDIA3*, *PPIB-ERP29* insertion at Sg7 |
| gRNA-302-CANXF | CTCAGTTTGCCATTATACCA | Target $P_{ScADH2}$ region in pre-integrated *CANX* cassette; |

| | | |
|---|---|---|
| | | used for *PDIA3* insertion upstream of *CANX* |
| gRNA-302-CANXR | TGGTATAATGGCAAACTGAG | Target $P_{ScADH2}$ region in pre-integrated *CANX* cassette; used for *PDIA3* insertion upstream of *CANX* |
| gRNA-302-CALRF | CAAGGAACACCTACATACCA | Target $P_{ScTEF2}$ region in pre-integrated *CALR* cassette; used for *PDIA3* insertion upstream of *CALR* |
| gRNA-302-CALRR | TGGTATGTAGGTGTTCCTTG | Target $P_{ScTEF2}$ region in pre-integrated *CALR* cassette; used for *PDIA3* insertion upstream of *CALR* |
| gRNA-302-2CF | AAAAATTCGCGTCTATACCA | Target $P_{ScPGK1}$ region in pre-integrated *PDIA3* cassette; used for *PPIB-ERP29* insertion upstream of *PDIA3* |
| gRNA-302-2CR | TGGTATAGACGCGAATTTTT | Target $P_{ScPGK1}$ region in pre-integrated *PDIA3* cassette; used for *PPIB-ERP29* insertion upstream of *PDIA3* |
| gRNA-702F | GATGAGCGAAACGAAACGTA | Target Sg19; used for *EDEM2* or *EDEM2-E117Q* insertion |

| | | |
|---|---|---|
| gRNA-702R | TACGTTTCGTTTCGCTCATC | Target Sg19; used for *EDEM2* or *EDEM2-E117Q* insertion |
| gRNA-GTB1-KO-F | TTTGAAATGGCCGGACCCTT | Target *GTB1* locus; used for *GTB1* deletion |
| gRNA-GTB1-KO-R | AAGGGTCCGGCCATTTCAAA | Target *GTB1* locus; used for *GTB1* deletion |
| gRNA-ROT2-KO-F | ACTTGAAGATCGGTGAAGAC | Target *ROT2* locus; used for *ROT2* deletion |
| gRNA-ROT2-KO-R | GTCTTCACCGATCTTCAAGT | Target *ROT2* locus; used for *ROT2* deletion |
| gRNA-MNS1-KO-F | GATGAGTTGAATTTAACCAC | Target *MNS1* locus; used for *MNS1* deletion |
| gRNA-MNS1-KO-R | GTGGTTAAATTCAACTCATC | Target *MNS1* locus; used for *MNS1* deletion |
| gRNA-CNE1-KO-F | GCTGTTGGTGCAACTTCAAC | Target *CNE1* locus; used for *CNE1* deletion |
| gRNA-CNE1-KO-R | GTTGAAGTTGCACCAACAGC | Target *CNE1* locus; used for *CNE1* deletion |
| gRNA-GTB1-KI-F | GAGCTGATTGTCATGCTACA | SpCas9 gRNA targeting the 3′ flanking region of *GTB1*; used for insertion of the *PRKCSH* cassette and *GTB1-E110A*, *GTB1-R629A*, or *GTB1-Y654F* cassettes. |

| | | |
|---|---|---|
| gRNA-GTB1-KI-R | TGTAGCATGACAATCAGCTC | SpCas9 gRNA targeting the 3′ flanking region of *GTB1*; used for insertion of the *PRKCSH* cassette and *GTB1-E110A*, *GTB1-R629A*, or *GTB1-Y654F* cassettes. |
| gRNA-ROT2-KI-F | ACCGGGACGGCGTATAACTG | SpCas9 gRNA targeting the 3′ flanking region of *ROT2*; used for insertion of the *GANAB* cassette. |
| gRNA-ROT2-KI-R | CAGTTATACGCCGTCCCGGT | SpCas9 gRNA targeting the 3′ flanking region of *ROT2*; used for insertion of the *GANAB* cassette. |
| UGGT1-donor-F | CTCCATTTCATTGATTATCTATGTG | Amplify Sg1 donor fragment for *UGGT1* or *UGGT1-D1358A* integration |
| UGGT1-donor-R | CGTACAAGTCCCCATTCCAGCCATA | Amplify Sg1 donor fragment for *UGGT1* or *UGGT1-D1358A* integration |
| UGGT2-donor-F | ACGAGGTGAGAAACGAATATTCCCT | Amplify Sg4 donor fragment for *UGGT2* or *UGGT2-D1333A* integration |
| UGGT2-donor-R | ATATTTATGCGTTGGCATTGGCTTA | Amplify Sg4 donor fragment for *UGGT2* or *UGGT2-D1333A* integration |

| | | |
|---|---|---|
| EDEM2-donor-F | TTTTTTCGGGTGTCGATGAGGATAT | Amplify Sg19 donor fragment for *EDEM2* or *EDEM2-E117Q* integration |
| EDEM2-donor-R | CCAAAAGTTTAACTCATGGGATCCC | Amplify Sg19 donor fragment for *EDEM2* or *EDEM2-E117Q* integration |
| 302-donor-F | CGCCGTGGATGAAGATGAAGGT | Amplify Sg7 donor fragment for *SEP15-U96C, CANX*, *CALR*, *PDIA3*, and *PPIB-ERP29* integration |
| 302-donor-R | AAGGAAAATGTCCATCCCTACAGG | Amplify Sg7 donor fragment for *SEP15-U96C, CANX*, *CALR*, *PDIA3*, and *PPIB-ERP29* integration |
| PDIA3-CANXup-donor-F | TTCCTTTCCTGTTCAATGCAGAATT | Amplify donor for inserting *PDIA3* upstream of pre-integrated *CANX* |
| PDIA3-CANXup-donor-R | CTGAAGACGAATTGGGAAGTATCGG | Amplify donor for inserting *PDIA3* upstream of pre-integrated *CANX* |
| PDIA3-CALRup-donor-F | TTCCTTTCCTGTTCAATGCAGAATT | Amplify donor for inserting *PDIA3* upstream of pre-integrated *CALR* |
| PDIA3-CALRup-donor-R | TAAATGTGTGGGTGCAACATGAATG | Amplify donor for inserting *PDIA3* upstream of pre-integrated *CALR* |

| | | |
|---|---|---|
| PPIB-ERP29-CNXp57-donor-F | CGCCGTGGATGAAGATGAAGGT | Amplify donor for inserting *PPIB-ERP29* in $KM_{CNX\text{-}p57}$ background |
| PPIB-ERP29-CNXp57-donor-R | TGTAAATGTAAGTTTCACGAGGTTC | Amplify donor for inserting *PPIB-ERP29* in $KM_{CNX\text{-}p57}$ or $KM_{CRT\text{-}p57}$ background |
| GTB1-locus-donor-F | AAGCGTCCAGCACGCCTCAG | Amplify donor for inserting E110A, R629A and Y654F mutation or PRKCSH into Km *GTB1* locus |
| GTB1-locus-donor-R | GGGCCAATACTATACCGACCACTG | Amplify donor for inserting E110A, R629A and Y654F mutation or PRKCSH into Km *GTB1* locus |
| ROT2-locus-donor-F | GCGGTCTATTTGGGCGTCAGTA | Amplify the *ROT2*-locus donor fragment from LHZ2051 for insertion of the *GANAB* cassette into the *ROT2* locus. |
| ROT2-locus-donor-R | GTATGGCTTCACCAGTAAGATCGCT | Amplify the *ROT2*-locus donor fragment from LHZ2051 for insertion of the *GANAB* cassette into the *ROT2* locus. |
| Human IgG Fc fragment coding sequence | gccgagccaaagtcttgtgacaaaacccacacttgtccaccttgtccagct ccagaattgttgggtggtccatctgtctttttgttcccaccaaagccaaagga caccttgatgatctcccgtacccccgaagtcacttgtgttgtcgttgacgtctc | This is not a primer but the codon-optimized coding sequence of the human IgG |

| | | |
|---|---|---|
| | tcacgaagatcccgaagttaaattcaactggtatgttgacggtgtcgaagtccacaacgccaagaccaagccacgtgaagagcagtacaactccacctacagagttgtctccgttttgaccgttttacaccaagattggttgaacggcaaggagtacaagtgtaaggtctccaacaaggctttgcccgctcctatcgagaagaccatttccaaggccaaaggccaaccaagagagccacaagtttacaccttgccaccttctagagacgagttgaccaagaatcaagtttccttgacttgtttggttaagggtttttatccatccgacatcgccgttgaatgggaatccaacggtcagcccgaaaacaactacaagaccaccccacccgttttggattctgacggttccttctttttgtactccaagttaactgttgataagtcccgttggcagcaaggtaacgtcttctcttgttccgtcatgcacgaggccttgcataaccactacactcaaaagtctttgtccttgtctccc | Fc fragment used to construct the secretory expression plasmid pExp-IgG Fc. |
| Sta2-Cas9 filler fragment | agtgacacgggttcgaatcccgtcagtgtcagcatgcaggtgcaggtccacctgcgtacgttttagtactctggaaacagaatctact | Synthetic DNA fragment used to construct the Sta2-Cas9 empty vector LHZ2065. This fragment contains two AarI sites for target-sequence insertion and the Sta2-Cas9 gRNA scaffold sequence. |
| Sta2-Cas9 gRNA scaffold sequence | gttttagtactctggaaacagaatctactaaaacaaggcaaaatgccgtgtttatctcgtcaacttgttggcgaga | Constant scaffold sequence contained within the Sta2-Cas9 filler fragment and required to form functional Sta2-Cas9 gRNAs after target-sequence insertion. |
